# The small ORFeome of *Tribolium castaneum* reveals deeply conserved small ORFs broadly expressed across tissues

**DOI:** 10.1101/2025.09.15.675444

**Authors:** Diego Guerra-Almeida, Mariana Freitas Nery, Jonathan Javier Mucherino-Muñoz, Evenilton Pessoa-Costa, Diogo Antonio Tschoeke, Rodrigo Nunes-da-Fonseca

## Abstract

Small open reading frames (small ORFs/smORFs/sORFs) were historically overlooked due to their short length (<100 codons) and detection challenges. It is widely accepted that small ORFs are rarely conserved across distant taxa, particularly in deeply rooted ancestors. Growing evidence supports their biological relevance and potential roles in gene innovation. Here, we integrated genomic, transcriptomic, and proteomic datasets to identify and characterize 454 putative small ORFs in *Tribolium castaneum*. Conservation analyses revealed 230 small ORFs conserved within Coleoptera, 167 in insects, 85 in arthropods, 39 in eukaryotes, and six in bacteria. Conserved small ORFs displayed higher GC content (∼44.87%), with bacterial-conserved sequences exceeding 50%. Functional enrichment revealed dozens of small ORFs associated with catalytic, transporter, structural, and regulatory roles. Binding site predictions indicated 254 small ORFs possess protein-protein interaction potential. Expression profiles across 14 RNA-seq libraries spanning multiple developmental stages showed 40.1% of small ORFs expressed (TPM >1) in at least one sample, and 16.5% expressed in five or more libraries. Conserved small ORFs showed broader and more stable expression, while lineage-specific ones displayed restricted expression, mostly in gonads or embryos. Isoformic and small CDSs were the most widely expressed classes. Notably, 29 small ORFs exhibited transcriptional and translational support (Swiss-Prot and/or Ribo-seq), 28 being small CDSs. A total of 390 small ORFs had potential paralogs, supporting gene duplication as a mechanism for small ORF emergence. Altogether, our findings reinforce that small ORFs, contrary to initial assumptions, can be deeply conserved and potentially perform key roles in evolutionary processes.

## INTRODUCTION

The advancement of large-scale DNA sequencing [1] has greatly contributed to our evolutionary understanding of life by providing extensive data for comparative analyses. In this context, several initiatives aim to sequence genomes across a wide variety of taxa. For instance, the i5K project is currently sequencing genomes of five thousand arthropod species [2], while the Genome 10K initiative seeks to sequence the genomes of at least one representative from each vertebrate genus [3]. In RNA science, deep transcriptome sequencing has drastically transformed molecular biology [4,5], particularly in the area of single-cell transcriptomics applied to evolutionary developmental biology [6]. Consequently, comparative studies focusing on previously poorly explored DNA sequences, such as “junk DNA”, have markedly increased in tandem with sequencing advances, opening new avenues for the discovery of novel genes and molecular innovations related to DNA expression [7,8]. In the last years, one of the most challenging components of junk DNA has gained recognition as a “hidden coding DNA world” [9,10]. These components are called small Open Reading Frames (small ORFs/smORFs/sORFs), which consist of small sequences potentially translated into peptides of less than 100 amino acids [10].

In the last years, dozens of small ORFs have been discovered as essential players in several biological contexts [11]. However, the reason small ORFs are referred to as a “hidden coding DNA world” is that millions of random and non-functional small ORFs are present in genomes by chance, due to their small size [12]. Consequently, ORFs shorter than 100 codons have historically been overlooked in annotation screenings [13]. Additionally, the assumption that small ORF peptides cannot fold into stable and functional structures has substantially contributed to their dismissal [14]. In fact, the discovery of coding and functional small ORFs is rare, making their computational detection among a myriad of random sequences a challenging task [15]. Most gene prediction software employs stringent parameters to evaluate potentially coding sequences; however, putative small ORFs often lack canonical features [16]. For instance, small ORF peptides commonly exhibit different codon-usage in comparison to canonical proteins [17], and may use alternative initiation codons[18]. Moreover, small ORF peptides can be translated from non-canonical mRNAs, such as circular RNAs (circRNAs) [19]. One promising strategy for assessing coding potential in small ORF studies involves evolutionary conservation analysis using the Basic Local Alignment Search Tool (BLAST) [20] and PhyloCSF [21]. However, deeply conserved small ORFs are usually overlooked in standard analysis, since point mutations result in more pronounced divergence in short sequence alignments compared to longer sequences. Consequently, putative small ORFs are likely to yield lower sequence similarity scores in conservation analysis [22]. Another long-established evolutionary strategy is the comparison between synonymous and non-synonymous changes, the Ka/Ks ratio. A Ka/Ks ratio below 1 indicates that a sequence is undergoing purifying selection, removing deleterious regions and preserving conserved zones [23]. Nonetheless, putative small ORFs may randomly achieve values below or above this threshold due to their small size [24]. In the last years, several programs and bioinformatic pipelines based on primary sequence analysis have been developed for small ORF prediction [25–28], albeit evidence suggests that selective pressure may act on the secondary structure of small ORF peptides independently of their amino acid sequence [29].

Experimental analyses also present significant challenges to the discovery of small ORFs. While studies on single small ORFs are crucial for identifying functional sequences, their application to whole genome analysis is limited by methodological, financial, and temporal constraints. Even advanced proteomic techniques often struggle to detect small ORF peptides, particularly those shorter than 20 amino acids known as dwarf small ORFs [30] as they are rapidly degraded by endogenous proteases [31]. Strategies based on ribosome profiling techniques (ribo-seq) offer advantages for small ORF detection; however, this approach cannot reliably distinguish functionally translated small ORFs from translational noise [32,33]. Additionally, the high cost and technical complexity of this method make this approach inaccessible to many research groups, particularly in low-income countries.

In this context, it is widely accepted that the discovery of new putative small ORFs is a challenging task that requires the combination of several complementary strategies. Thus far, genome-wide bioinformatic identification of small ORFs in the Arthropoda phylum has only been performed in the fruit fly *Drosophila melanogaster* [24]. The Arthropoda phylum (including insects, crustaceans, spiders, ticks, scorpions, and others) is the most successful metazoan clade, representing approximately 85% of all described animals [34], thus it is important to extend small ORF analysis to other groups.

In this study, we perform a low-cost bioinformatic approach to predict and evaluate the evolution of small ORFs in the red flour beetle *T. castaneum*. The red flour beetle was selected as the reference taxon in this study for several reasons. First, it has a fully sequenced and well-annotated genome, including recent long-read assemblies [35]. Second, it offers a robust set of genetic tools, including systemic RNA interference and efficient CRISPR-based genome editing [36–38], which facilitate the functional investigation of newly identified small ORF genes. Third, *T. castaneum* exhibits a more conserved gene repertoire compared to more recently evolved insect models.

Our findings offer new insights into the origin, conservation, and potential functions of small ORFs, establishing a foundation for future experimental validation in this promising model for small ORF research.

## RESULTS

### Putative Small ORFs Discovered in *T. castaneum*

The *T. castaneum* intergenic DNA spans 223.5 Mb (megabases). Approximately 1.5 million small ORFs were detected across both DNA strands, resulting in a density of one small ORF per 147 base pairs (bp).

Given that evolutionary conservation is an important hint of functionality, we applied a comparative genomic filter to identify conserved small ORFs within the pool of *de novo* predicted sequences. This analysis was concentrated on four coleopteran species with publicly available genomes at the time of analysis. Our results reveal that 14,878 small ORFs are conserved across these coleopteran species. This number is 100 times lower than the amount of *de novo* predicted small ORFs, which suggests that millions of random sequences were excluded (Figure 1A).

**Figure 1.**
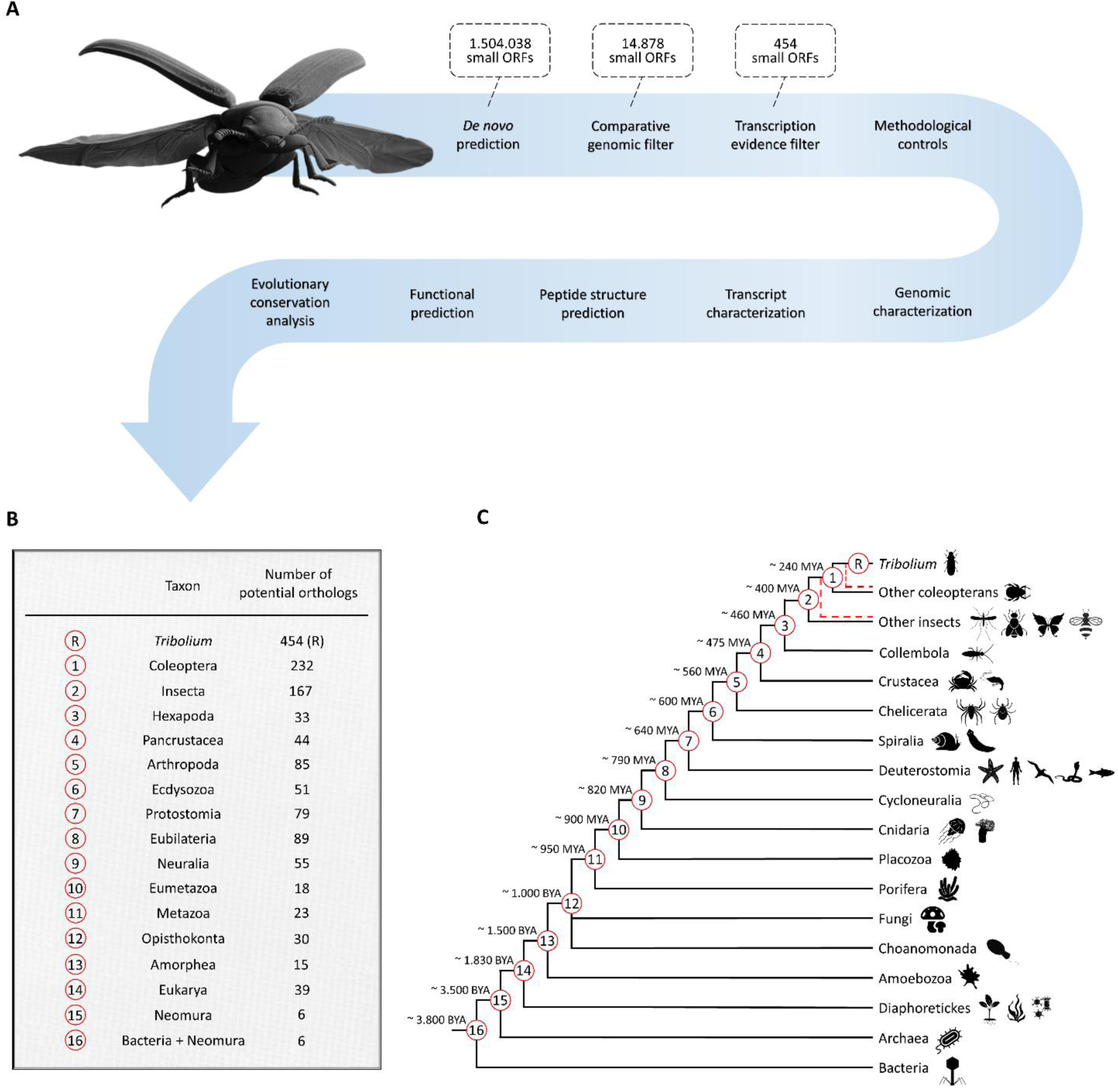
Workflow for the Prediction and Evolutionary Analysis of Small ORFs in *T. castaneum*. **(A)** Schematic overview of the methodological steps used for the detection and characterization of small ORFs, including both positive and negative controls, as well as the evolutionary analysis. **(B)** Distribution of potential orthologous small ORFs conserved across multiple taxa. Numerical labels to the left of each taxon correspond to ancestral nodes referenced in panel C. The number of species with available RefSeq transcripts used in the analysis is shown to the right of each taxon name. **(C)** Phylogenetic tree with *T. castaneum* as the reference taxon (marked “R”). Red dashed lines indicate that species within a taxon may also derive from the preceding branch. Each branch represents a clade sharing a common ancestor with *T. castaneum*, indicating the taxonomic range in which potential small ORF orthologs were identified. Phylogenetic tree based on [82,102,105,124].

To prevent the filtering of redundant sequences from potential assembly errors or recently diverged genes, we applied an additional redundancy filter (≥ 80% identity), thereby excluding 1,221 potentially redundant small ORFs while retaining the largest representatives. This additional step reduced the total number of coleopteran-conserved small ORFs to 13,657 sequences with an average size of 55 codons.

Finally, we compared the coleopteran-conserved small ORFs with the complete set of small ORFs in *T. castaneum* RefSeq transcripts available on NCBI. Importantly, the use of RefSeq transcripts in this filter is not conclusive in ensuring gene expression. Although the RefSeq database is a comprehensive, non-redundant, and well-annotated collection of reference sequences from NCBI, it also includes transcripts predicted by computational methods in cases where experimental evidence is limited or absent. Thereby, this comparison identified 454 small ORFs, a number approximately 30 times lower than the amount of small ORFs filtered in the previous step (Figure 1A). The shortest putative small ORF detected was 12 codons, a size rarely reported in the literature; their average size is 58 codons (Figure 2A). This refined subset constitutes the putative small ORFs of *T. castaneum,* hereafter named *T. castaneum* small ORFeome.

**Figure 2.**
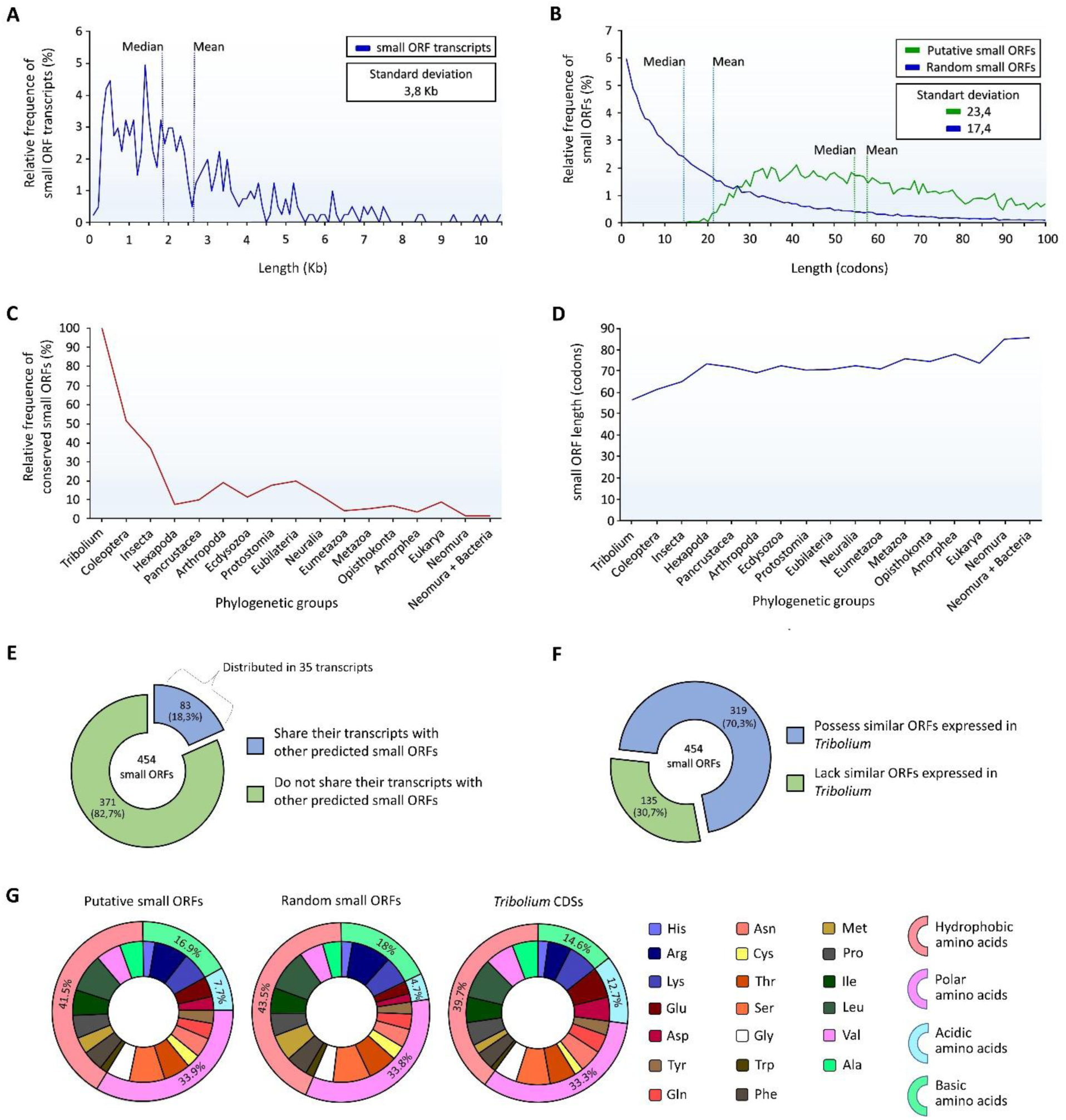
Genomic Characterization of Putative Small ORFs in *T. castaneum*. **(A)** Relative distribution of putative small ORF transcript lengths. Median = 1.9 kb; mean = 2.6 kb. **(B)** Relative frequency of lengths (in codons) for putative versus random small ORFs in *T. castaneum*. **(C)** Frequency of small ORFs with putative orthologs in each evolutionary group, relative to the total of 454 predicted small ORFs. **(D)** Average length of putative small ORFs in *T. castaneum* across different levels of evolutionary conservation. **(E)** Mapping of putative small ORFs that co-occur with other predicted small ORFs within the same transcript. **(F)** Proportion of small ORFs with or without similar ORFs (putative paralogs) of any size expressed in *T. castaneum*. **(G)** Amino acid composition per 1,000 residues in predicted peptides from *T. castaneum* small ORFs, categorized by electrostatic properties. Amino acid abbreviations: Ala (alanine), Arg (arginine), Asn (asparagine), Asp (aspartic acid), Cys (cysteine), Gln (glutamine), Glu (glutamic acid), Gly (glycine), His (histidine), Ile (isoleucine), Leu (leucine), Lys (lysine), Met (methionine), Phe (phenylalanine), Pro (proline), Ser (serine), Thr (threonine), Trp (tryptophan), Tyr (tyrosine), and Val (valine).

### Negative Control Analysis Highlights the Specificity of the Small ORF Identification Method

To evaluate the stringency of the filters against known random sequences, a negative control was conducted using small ORFs extracted from the reverse and non-complementary genome sequence of *T. castaneum*. These random sequences were subjected to the same prediction filters applied to putative small ORFs. The *de novo* detection step predicted over two million small ORFs in the reverse and non-complementary genome sequence, which is expected as small ORFs are conceptually merely DNA stretches containing a start and stop codon within the same frame.

Only 95 random small ORFs were obtained during the comparative genomics filtering. From these, 93 were predominantly composed of tandem repeat regions, showing high similarity to sequences of the same type in other coleopterans. Repeat regions are low-complexity sequences, which can lead to false-positive results in similarity search tools such as BLAST.

No random small ORFs were obtained during the RefSeq transcript filtering step, using an E-value threshold of 0.001. This search was repeated using less stringent E-values of 0.01 and 0.1. These values were previously applied for searches against RefSeq transcripts from distant eukaryotic and prokaryotic groups, respectively. However, no random small ORFs were obtained. Finally, the analysis was repeated using even higher E-values. Only when the E-value was set to 10 did the searches return six false-positive results. Among these, two sequences exhibited 90% similarity, although all bitscores remained consistently below 27. Therefore, using the E-value as a quality indicator is more reliable than relying merely on similarity scores, since even random small ORFs can achieve high similarity levels. Thereby, dwarf small ORFs with diminutive functional motifs could easily be overlooked in similarity searches. In conclusion, the filtering methodology adopted here is stringent against known random small ORFs.

### Evolutionary Analysis of *T. castaneum* Small ORFs Along the Tree of Life

Given the relevance of evolutionary conservation in identifying potentially functional DNA regions, we compared the putative small ORFs with those extracted from RefSeq transcripts of 15 distantly related groups, including prokaryotic clades (Figure 1B). Our analysis revealed that dozens of putatively functional small ORFs exhibit high levels of conservation. Approximately 230 potential orthologs were identified in other coleopterans, with at least 167 conserved in the common ancestor of insects. This number decreases to about 85 sequences or fewer outside Insecta in other arthropods, and further declines to 39 conserved small ORFs in the common ancestor of Eukarya. Notably, our analysis suggests that six putative small ORFs may have orthologs in Bacteria (Figure 1C).

The number of conserved small ORFs fluctuated across taxonomic groups, reflecting differences in the RefSeq database content for each taxon. For instance, we identified 33 potential orthologs in Hexapoda, while 89 conserved sequences were detected in the more distantly related Eubilateria group (Figure 1B). Overall, the total number of conserved sequences tends to decline with increasing evolutionary distance (Figure 2C), while the average small ORF length tends to increase, ranging from 70 to 80 codons in distant ancestor groups (Figure 2D).

Regarding the analysis of taxonomically restricted small ORFs, 163 sequences likely emerged in *T. castaneum* lineage, followed by 80 small ORFs potentially emerging in Coleoptera, 63 in Insecta, 31 in Eubilateria and 19 in Neuralia. Moving further, Eukarya has 31 potentially restricted sequences, followed by Neomura and Bacteria (Last Universal Common Ancestor – LUCA) with 4 and 6 sequences, respectively (Additional file 1). Importantly, although these inferences are based on the taxa where the transcription evidence emerged, they may be non-expressed in the genomes of other groups. For example, all small ORFs restricted to *T. castaneum* show genomic-level conservation with other beetles, as shown by the comparative genomics filter.

### Genomic Characterization Shows Differential Distribution of Putative Small ORFs Between Autosomal and Sex Chromosomes

The ratio of chromosome length to the emergence of putative small ORFs indicates the amount of DNA sequence required for the evolution of a unique small ORF. Chromosomes 3 and 10 exhibit higher ratios, suggesting that more extensive DNA sequences are needed for the development of putative small ORFs on these chromosomes. In contrast, the sex chromosomes X and Y generate more putative small ORFs per stretch of DNA (Table 1).

**Table 1.**
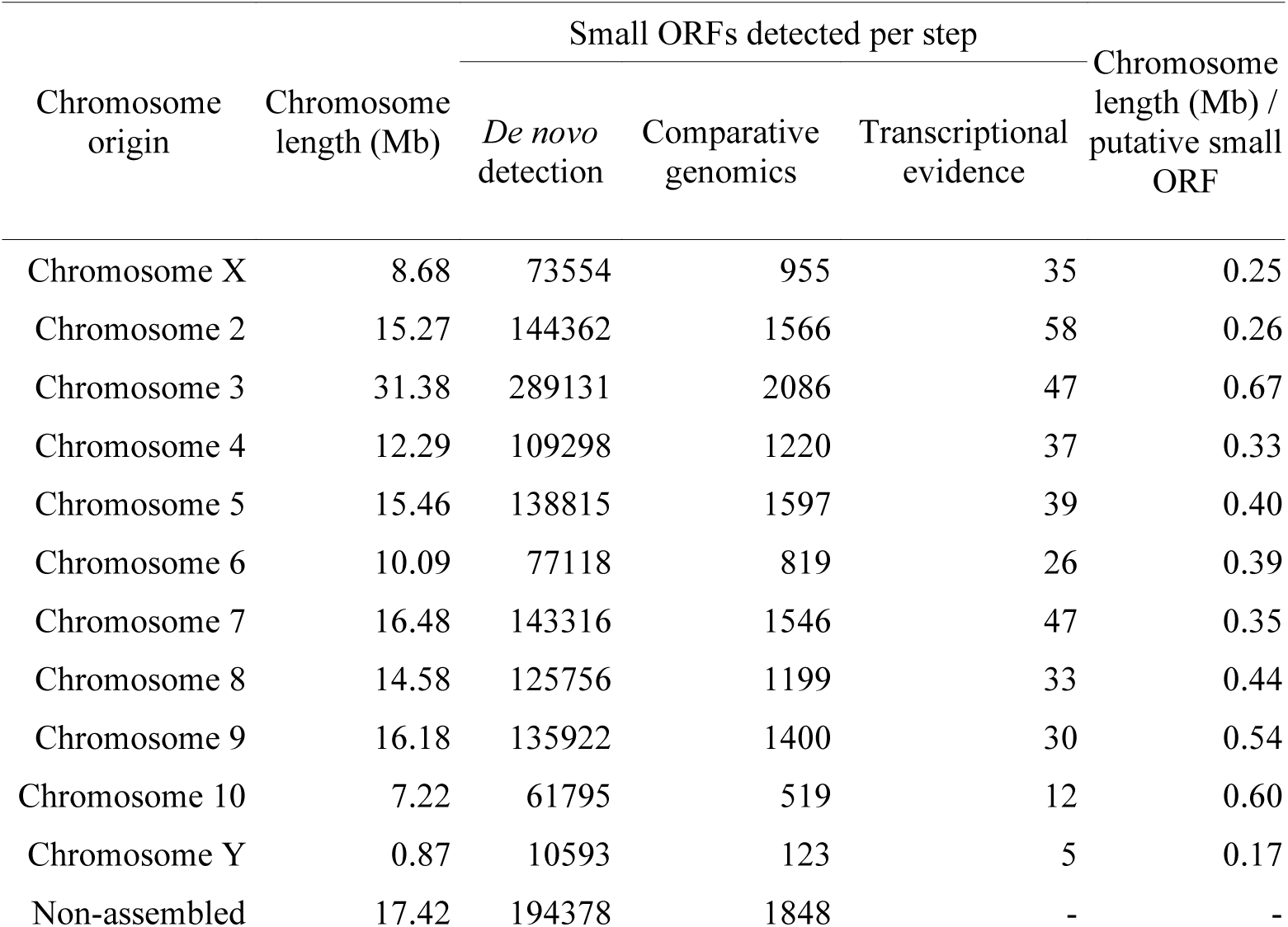
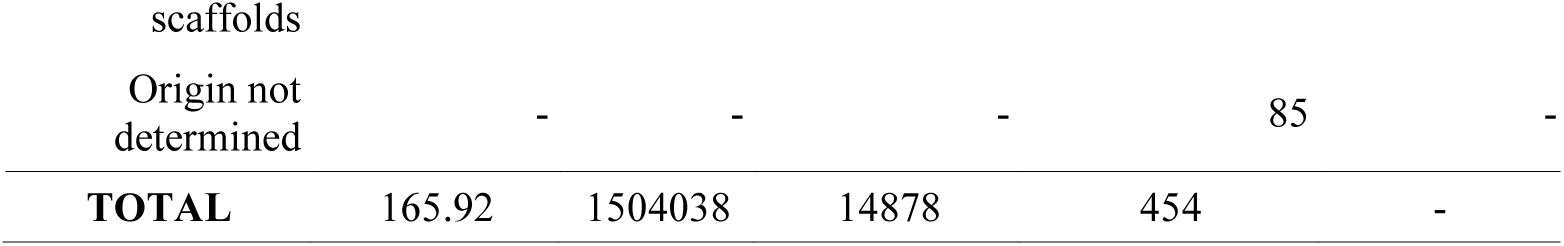
Number of small ORFs per chromosome and ratio of chromosome length to putative small ORF. . The 85 small ORFs from the transcriptional evidence filter with undetermined chromosomal origin correspond to transcripts whose chromosomal origin is undefined.

### Small ORF Transcript Analysis

Previous study indicates that functional metazoan small ORFs are typically found in transcripts ranging from 400 to 600 bp in length [39]. In *T. castaneum*, small ORF transcripts have an average size of approximately 2.6 kb. However, the median transcript size is 1.9 kb, highlighting the high frequency of transcripts within the 500 bp to 2.3 kb range (Figure 2B). Remarkably, the two largest small ORF transcripts are 27 kb and 63 kb in length, while the smallest ones measure 90 bp and 160 bp.

Considering that the well-characterized small ORF gene *mille-pattes*/*tarsal-less*/*polished-rice* is known to exhibit a polycistronic transcript organization [40–44] we explored the potential for similar arrangements in *T. castaneum*. Our analysis identified 35 potential polycistronic genes, each containing two or more putative small ORFs. These 35 genes encompass 83 (18%) of the detected putative small ORFs (Figure 2E). Notably, while genes shared by both small and large coding ORFs are conceptually recognized as polycistronic RNAs [45], our analysis focused on transcripts containing strictly small ORFs, since small ORFs sharing mRNAs with large ORFs are more appropriately classified as alternative small ORFs [10].

Furthermore, we searched for potential paralogs of any size within *T. castaneum* RefSeq transcripts. Our results show that 70% of putative small ORFs have one or more similar ORFs (Figure 2F). Among these potential paralogs, 153 are large ORFs, and 237 are small ORFs.

### GC Content and Amino Acid Usage Analysis

Our data show that the 454 putative small ORFs in *T. castaneum* have a GC content (∼44.87%) closely matching that of random small ORFs (∼43.72%) (Additional file, table 3). Hence, while these small ORFs do not appear drastically different from random small ORFs in compositional terms, they do stand out relative to the overall genomic background, suggesting that some may be under constraints typical of coding sequences [46,47]. Notably, the small ORFs conserved across prokaryotes exceed 50% GC (Additional file, table 4). Additionally, codon-position analysis indicates that GC levels are lowest at the second site and highest at the third, a pattern also observed in the annotated CDSs of *T. castaneum* (Additional file, table 3).

The amino acid usage of the putative small ORFs of *T. castaneum* is more similar to random sequences than to canonical CDSs (Figure 2G). Our analysis reveals that the frequency of arginine in putative small ORFs is higher in comparison to canonical CDSs. Interestingly, random small ORFs exhibit an even higher frequency of arginine compared to non-random sequences (Figure 2G), a phenomenon that warrants further investigation (Additional file 2, table 5).

The putative small ORFs of *T. castaneum* contain eight amino acids with frequencies comparable to those reported in α-helix regions [48], such as, alanine (Ala), methionine (Met), phenylalanine (Phe), cysteine (Cys), glutamine (Gln), tyrosine (Tyr), aspartic acid (Asp), and glutamic acid (Glu). Furthermore, these small ORFs exhibit a notable prevalence of hydrophobic amino acids, a critical feature for transmembrane peptides [49,50].

### Small ORF Classes Display Different Patterns

We manually categorized the putative small ORFs following the transcription feature classification proposed by Guerra-Almeida et al. (2021), which classifies small ORFs as either non-expressed (intergenic) or expressed (genic). Within the expressed category, genic small ORFs are further subdivided based on their transcriptional context. Thereby, genic small ORFs are located either in non-coding RNAs (ncRNAs) or canonical mRNAs. Small ORFs in ncRNAs are further distinguished by their presence in either small or long RNA molecules (Figure 3).

**Figure 3.**
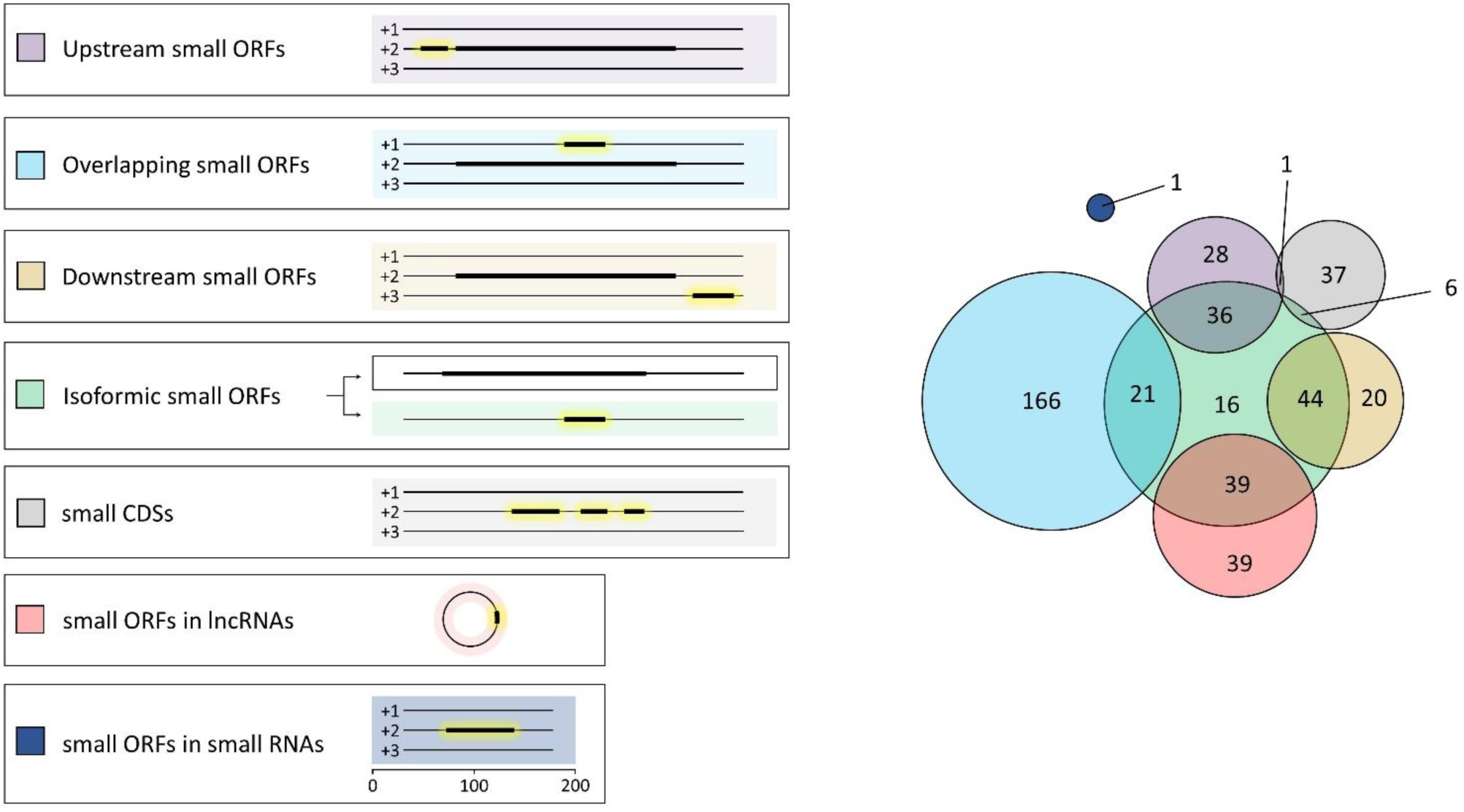
Classification of Putative Small ORFs in *T. castaneum*. **Left panel**: Color-coded legend representing each transcriptional class of small ORFs, along with schematic examples illustrating the transcriptional features associated with each class. In each transcript, small ORFs are highlighted in yellow. **Right panel**: Venn diagram showing the distribution of putative small ORFs across distinct transcriptional classes. Notably, individual small ORFs may belong to multiple classes based on their transcriptional context. The classification framework is adapted from Guerra-Almeida et al. (2021) [10].

On the other hand, small ORFs in canonical mRNAs are classified into several specific classes: alternative small ORFs (upstream, overlapping, or downstream to a canonical CDS in the RNA), isoformic small ORFs (isoforms derived from large CDSs), and small CDS (known coding small ORFs in canonical-like transcripts) (Figure 3).

In our dataset, the distribution of small ORF categories highlights notable patterns in their representation. The most frequent group is overlapping small ORFs, appearing 187 times, followed by isoformic small ORFs with 162 instances, and small ORFs found in long non-coding RNAs (lncRNAs), which occur 78 times. Classes with moderate representation include upstream small ORFs (65 entries), downstream small ORFs (64 entries), and small CDS with 44 recorded cases. The rarest category is small ORFs within small RNAs, observed only once (Figure 3).

The evolutionary origins of small ORFs vary significantly across classes. From the 129 downstream and upstream small ORFs, 83 emerged in more recent common ancestors, such as *T. castaneum* and other insect lineages, suggesting adaptive roles specific to these groups. In contrast, among the 44 small CDS 38 are linked to more ancient ancestors, including broad eukaryotic lineages starting from Neuralia and even Bacteria, reflecting potential highly conserved functions (Additional file 1).

Analysis of molecular function reveals additional insights. For example, 16 isoformic small ORFs are associated with binding activities and 12 with catalytic roles. Regarding small CDSs, nine are associated with mitochondrial bioenergetics, whereas another nine are linked to gene-expression processes, such as RNA processing, transcription, and translation. Classes like overlapping small ORFs and downstream small ORFs have a substantial number of sequences without predicted protein family associations, indicating potential functional specialization or non-annotated roles (Additional file 1).

### Predicted Secondary Structures, Protein Binding Site and Functional enrichment

254 small ORF-predicted peptides exhibit predicted protein binding sites (Additional file 1). Sequences exhibiting mixed secondary structure elements, such as helices and sheets, are more likely to include annotated binding sites, often involving N-terminal residues, highlighting the potential importance of these regions in interaction dynamics. 47 sequences display extended binding sites longer than 10 amino acids (Additional file 1). These extended regions may represent molecules with increased specificity or stability in protein-protein interactions. In contrast, sequences with highly repetitive coil structures, which are the most frequent motifs found, are predominantly associated with a lack of binding sites, suggesting disordered or transiently interacting domains [51].

Furthermore, sequences with annotated binding sites exhibit associations with specific biological processes, including protein glycosylation and translation, underscoring their potential involvement in essential cellular functions (Additional file 1). These entries are also linked to zinc ion binding and metalloendopeptidase activity, highlighting their putative roles in enzymatic catalysis and regulatory mechanisms [52,53]. Notable protein family associations further emphasize their functional diversity. For instance, a sequence predicted to encode a cytochrome C oxidase biogenesis protein Cmc1-like is linked to the binding site D82-L84, suggesting a role in mitochondrial assembly [54]. Similarly, a sequence associated with a potential dolichol-phosphate mannosyltransferase subunit 3, mapped to the binding site E6-V10 / A13-A16, points to a putative role in glycoprotein biosynthesis [55], reflecting the potential involvement of these small ORFs in complex biochemical pathways (Additional file 1).

Regarding the 47 small ORF-predicted peptides with protein binding sites longer than 10 amino acids, 34 sequences lack clear protein family assignments, indicating a high degree of under-characterization (Additional file 1). However, notable examples include a sequence associated with ecdysis-triggering hormone activity and another linked to the adipokinetic hormone/red pigment-concentrating hormone family, both of which suggest specialized roles in hormonal regulation, such as molting and energy mobilization [56,57]. Additional associations highlight broader functional diversity, including a sequence linked to the RNA polymerase archaeal subunit P/eukaryotic subunit RPABC4, which points to potential involvement in transcriptional processes [58], and another entry associated with pyruvate carboxylase, indicative of participation in central metabolic pathways [59]. Furthermore, a sequence mapped to the Sec61 translocon complex underscores its structural and possibly functional role in protein translocation across cellular membranes [60] (Additional file 1).

A subset of 45 small ORF-encoded peptide sequences contain protein family annotations (Additional file 1). Mitochondrial bioenergetics represents the most prevalent functional category: ten small ORFs encode subunits or assembly/biogenesis factors of oxidative phosphorylation complexes I, III, IV, and V. Membrane signaling and ion homeostasis constitute the second major group, comprising six sequences that correspond to transmembrane chemoreceptors, epithelial sodium channels, or other membrane proteins. Components of intracellular trafficking pathways, including TOM7, Sec61-β, BBSome-10, and LITAF, account for a further 7%, while regulators of gene expression (ribosomal proteins, RNA polymerase subunits, THAP DNA-binding factors, and splicing and chromatin modifiers) collectively comprise approximately 15%. Finally, a small set of neuropeptide hormones and stress-response proteins highlights the functional breadth of small ORFs, suggesting roles that span energy production, signaling, and cellular homeostasis.

393 small ORFs lack any detected gene ontology data. Among the remaining 64 sequences, several annotations highlight their potential functional and structural roles. Among the retrieved biological-process GO terms, proteolytic pathways were the most frequent (7 terms), together with transport-related processes (7 terms), encompassing ion movement, proton translocation, and protein import into mitochondria. DNA transposition or integration appeared six times, while central carbon metabolic routes, including carbohydrate, 2-oxoglutarate, pyruvate metabolism, and gluconeogenesis, accounted for four terms. Functions related to RNA processing and export comprised three entries, matching the three terms assigned to sensory perception of taste. Energy-related processes were represented by two terms linked to mitochondrial electron transport or ATP synthesis, whereas gene-expression steps (translation and basal transcription) contributed four terms. Smaller categories included cell signaling mediated by small GTPases or neuropeptides (2 terms), cilium assembly (1 term), and protein glycosylation (1 term), underscoring the functional breadth encompassed by the annotated set.

Among the 53 molecular-function GO terms found (Additional file 1), binding activities predominate (25 terms), spanning nucleic-acid, protein, metal-ion and nucleotide interactions. Catalytic functions appear 14 times, driven largely by proteases, transposases and metabolic enzymes. Five entries describe enzyme regulators, including guanyl-nucleotide exchange factors, kinase regulators and serine-type endopeptidase inhibitors. Transport and channel activities are represented by three terms related to sodium and proton conduction, while structural roles (ribosomal and chromatin components) and hormone-related signaling functions each contribute three terms. Collectively, the annotated set is enriched in binding and catalytic capabilities, yet still exhibits notable diversity in regulation, membrane transport, structural support and endocrine signaling.

Of the 25 cellular-component annotations retrieved (Additional file 1), mitochondrial structures are the most represented (7 terms), spanning respiratory-chain complex III, the F(o) sector of ATP synthase, oxoglutarate dehydrogenase, the outer-membrane import translocase and the organelle itself. Generic membrane assignments appear five times, highlighting peptides broadly associated with lipid bilayers. Five annotations map to nuclear or chromatin-associated assemblies, two nucleosome hits plus the Set1C/COMPASS and SAGA histone-modifying complexes and a nuclear pore entry. The dataset also includes two ribosomal terms and two trafficking complexes (BBSome and the Sec61 translocon). Single entries correspond to the V-type ATPase proton pump, the endoplasmic reticulum, the extracellular region and the anaphase-promoting complex, together indicating a diverse cellular distribution that nonetheless converges on mitochondria, membranes and chromatin-related compartments.

### Expression Analysis of small ORF Candidate Genes Across Developmental Stages and Tissues in *T. castaneum*

Expression levels of the 454 predicted small ORFs in 14 RNA-seq libraries of *T. castaneum* were normalized as transcripts per million (TPM) and visualized using a circular Circos plot (Figure 4). A total of 40.1% (182 small ORFs) were expressed in at least one tissue (TPM > 1), while 16.5% (75 small ORFs) were expressed in five or more libraries, suggesting moderate pleiotropy (Additional file 1). The remaining 59.9% displayed undetectable or minimal expression under the analyzed experimental conditions (Additional file 1). Small CDSs (n = 44) were among the most consistently expressed categories, with 61.3% (27/44) showing detectable expression and 40.9% (18/44) expressed in multiple tissues. Isoformic small ORFs (n = 162) showed intermediate expression levels, with 54.9% (89/162) expressed in at least one tissue and 27.2% (44/162) detected in five or more libraries. Overlapping small ORFs (n = 187), despite being the most numerous class, exhibited more restricted expression patterns: 38.5% (72/187) were expressed, and only 11.2% (21/187) were detected in five or more tissues. lncRNA-associated small ORFs (n = 78) showed high tissue specificity, with 30.7% (24/78) expressed predominantly in ovaries and embryos. Upstream (uORFs; n = 65) and downstream small ORFs (n = 64) displayed similar trends, with 35.4% (23/65) and 34.4% (22/64) expressed, respectively, mainly in testes and embryos.

**Figure 4.**
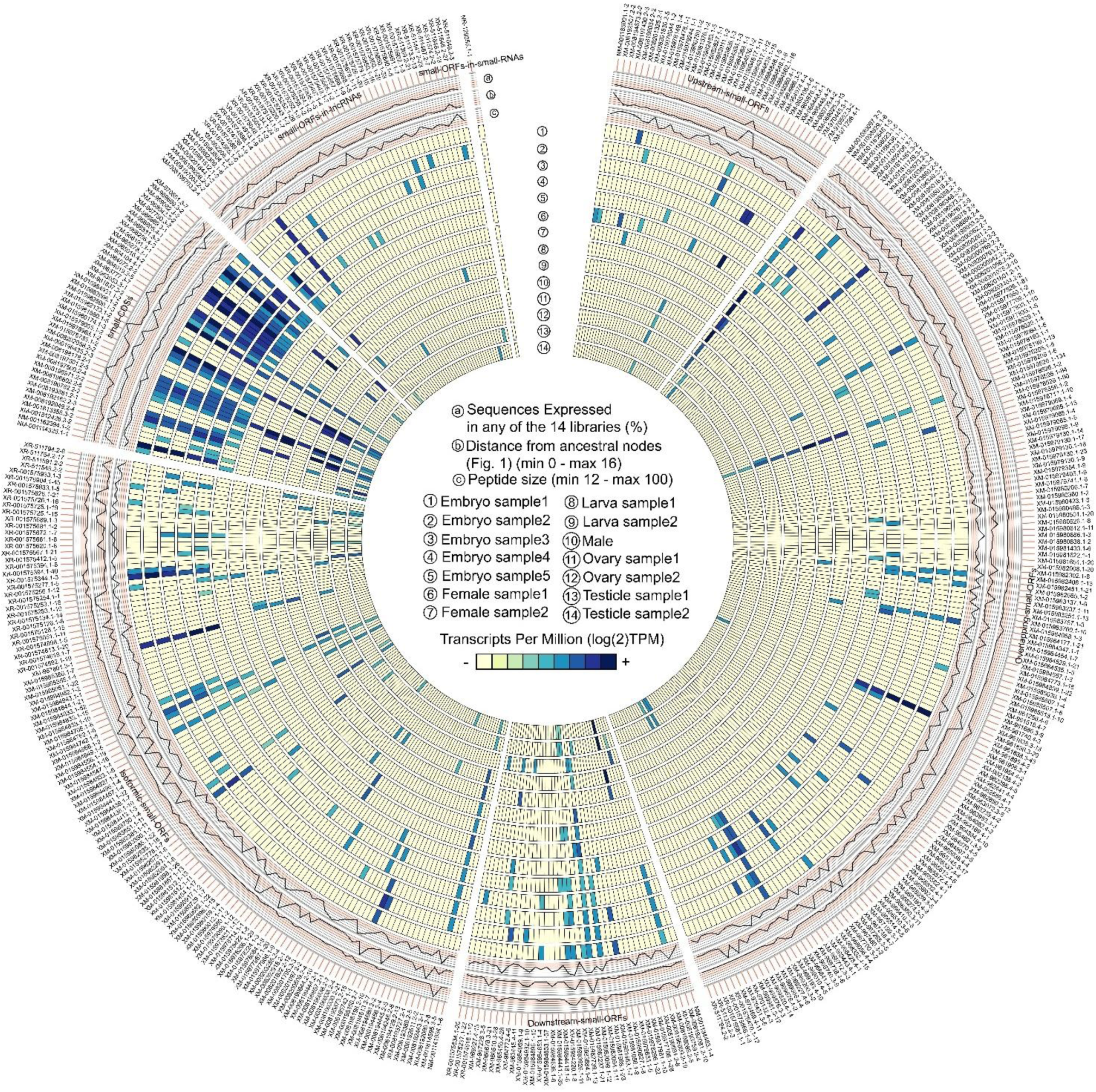
Expression profiles of predicted small ORFs across developmental stages and tissues in *T. castaneum*. Circular heatmap showing the expression levels (log₂TPM) of 454 predicted small ORFs across 14 RNA-seq libraries, including embryos (stages 1–5), larvae (samples 1–2), adult females (samples 1–2), adult males, ovaries (samples 1–2), and testes (samples 1–2). The outermost ring categorizes small ORFs by transcriptional class: overlapping, upstream, downstream, isoformic, small CDSs, small ORFs in lncRNAs, and small ORFs in small RNAs. Inner concentric rings represent expression values per library, with color intensity ranging from light yellow (low or undetectable expression) to dark blue (high expression). Additional data tracks include: (a) proportion of libraries in which the small ORF is expressed, (b) evolutionary conservation (relative to ancestral divergence), and (c) predicted peptide length. The figure reveals distinct tissue- and stage-specific expression patterns. Conserved small ORFs tend to be broadly expressed across samples, while lineage-specific small ORFs show more restricted expression, often with high expression levels in embryonic, gonadal, or larval tissues.

These class-specific differences were also reflected in average TPM levels, with isoformic small ORFs and small CDSs exhibiting broader and more stable expression across tissues, whereas overlapping, upstream/downstream, and lncRNA-associated small ORFs presented more restricted expression profiles (Figure 4 and Additional file 1). Evolutionary analyses showed that Eukarya- and Metazoa-conserved small ORFs exhibited significantly broader tissue expression and higher pleiotropy compared to lineage-specific ORFs. In contrast, *Tribolium* and Coleoptera-specific small ORFs were more narrowly expressed, typically restricted to one or two tissues, most often gonads or early embryos. Notably, embryonic samples contained the highest number of expressed small ORFs (n = 135), suggesting conserved roles in development, while testes exhibited the highest number of tissue-specific small ORFs (n = 52), supporting their role as genomic niches for novel regulatory peptides.

Interestingly, clusters of small ORFs located within the same transcript, suggesting potential polycistronic organization, showed coordinated expression patterns, indicating possible co-regulation. Collectively, these results demonstrate that a substantial fraction of the predicted small ORFs are transcriptionally active in vivo, supporting their biological relevance and providing a foundation for future functional characterization in *T. castaneum*.

### At least 40 Small ORFs Display Experimental Evidence of Translation

Small ORFs with experimental evidence of translation on ribo-seq and Swiss-Prot databases were assessed, composing a subset of small ORFs with strong translational support and functional potential. This analysis also serves as a positive validation of the methodology. Among this dataset, eight sequences exhibit ribo-seq evidence, while 33 small ORFs are supported by Swiss-Prot annotations. Importantly, six entries display evidence from both sources (Additional file 1).

Cross-referencing small ORFs with experimental translation evidence (Swiss-Prot and/or Ribo-Seq) and RNA-seq data identified 29 small ORFs that were both translated and transcriptionally active (TPM > 1 in at least one tissue-specific library). Of these, the vast majority (28) were classified as small CDSs, while a single upstream small ORF (NM_001185001.1_2) was also detected. This strong enrichment for small CDSs class among the translated and transcribed small ORFs supports the idea that these ORFs are not only translated but also transcriptionally integrated into functional developmental programs, particularly during embryogenesis. Notably, no lncRNA-associated, overlapping, or downstream small ORFs were detected in the translated and expressed subset, suggesting that these classes may be less likely to undergo stable translation or are translated in tissues not yet analyzed.

The average length of ribo-seq-supported sequences is approximately 76 amino acids, while those supported by Swiss-Prot are slightly longer, averaging 80 amino acids. For sequences with overlapping evidence from both datasets, the average length increases to 82 amino acids.

The analysis of ancestral origins highlights interesting results. For example, five sequences with ribo-seq evidence trace back to Eukarya origin, reflecting their broad evolutionary conservation. Two sequences potentially originate in more recent groups, such as Amorphea and Metazoa, respectively, while one entry points to Bacteria origin. In sequences supported by Swiss-Prot, Eukarya also remains the most common origin, with 11 occurrences, although other groups such as Neomura (4), Neuralia (4), Opisthokonta (3) and Metazoa (3) are also well represented, with a smaller number of entries tracing back to Arthropoda, Eubilateria, and Bacteria (Additional file 1). These results indicate that sequences with translational evidence tend to be longer and are often conserved across diverse evolutionary lineages, particularly within eukaryotes.

Regarding functional annotation, Swiss-Prot and ribo-seq-supported sequences are enriched for potential roles in mitochondrial processes and RNA splicing (Additional file 1). For example, families such as cytochrome B-C1 complex subunits and cytochrome C oxidase assembly protein PET191 are linked to mitochondrial electron transport and respiratory functions [61,62], while the splicing factor 3B subunit 5 family suggests involvement in mRNA processing [63]. Additionally, dolichol-phosphate mannosyltransferase subunit 3, associated with glycoprotein biosynthesis [55], reflects functional diversity within this subset. Furthermore, protein families like mitochondrial import receptor subunit TOM7, indicative of protein translocation [64], and hormonal regulators such as the eclosion hormone and adipokinetic hormone/red pigment-concentrating hormone families, which points to potential roles in molting and energy mobilization [56,57], expand this diversity. Notably, sequences with translation evidence from both ribo-seq and Swiss-Prot databases reinforce these findings, particularly for families involved in mitochondrial energy metabolism, RNA splicing, and protein transport, such as the cytochrome B-C1 complex and splicing factor 3B subunit 5 (Additional file 1).

## DISCUSSION

Several studies on small ORF prediction employ strategies based on transcriptomic and proteomic analyses [51,65–69]. However, approaches that integrate genomic data offer a relevant perspective, as they enable the identification of conserved genes that are functional in any tissue-specific expression, stage-specific developmental patterns, as well as other intrinsic and extrinsic factors [70]. For this reason, the integrative approach performed here aims to provide an in-depth analysis, bridging the three pillars of the molecular paradigm—genome, transcriptome, and proteome—through an evolutionary perspective, since evolutionary conservation is an accepted indicator of functionality [71].

The number of 454 predicted small ORFs correspond to approximately 2% of the amount of previously annotated proteins in the *T. castaneum* RefSeq database. Interestingly, this number closely resembles the 401 small ORFs identified in a similar study conducted on *Drosophila melanogaster* [24]. The ratio of the total genome size (in Mb) to the number of small ORFs identified (in units) is 0.36 in the beetle and 0.34 in the fruit fly, indicating that the stringency levels between the two methodologies are similar.

### Chromosomes and Their Small ORF Distribution

Chromosomes 3 and 10 are the least efficient at generating small ORFs, as they require more DNA extension per small ORF compared to other chromosomes (Table 1). This result might be explained by the higher density of satellite DNA within euchromatic regions found in chromosomes 3 and 10 in comparison to other autosomes in the beetle [72]. Satellite DNA, composed of tandemly repetitive sequences, creates an unfavorable environment for the emergence of coding sequences, thereby hindering the evolution of novel small ORFs on these chromosomes [73].

Sex chromosomes are characterized by unique ploidy and inheritance patterns that support essential roles in numerous biological processes, including sex determination, epigenetics, gene expression regulation, and genome organization in insects [74]. While the interplay between small ORFs and these distinctive roles is not yet explored, our findings suggest that the X and Y chromosomes are the most efficient at generating small ORFs (Table 1). The X chromosome has fewer repetitive regions within euchromatin compared to other chromosomes and displays a high rate of mutation and adaptive evolution, likely influenced by its hemizygous state in males [72,74].

In contrast, the Y chromosome in *T. castaneum* is highly degenerated and compact, consisting predominantly of its centromeric region and extensive stretches of heterochromatin [74,75]. The Y chromosome is also non-recombining, evolves at a slow rate, and contains very few functional genes. In the suborder Polyphaga, which includes *T. castaneum*, approximately 8% of males completely lack the Y chromosome (X0 males) without any noticeable phenotypic or reproductive changes; this phenomenon reaches 100% in certain beetle groups [76]. However, in *T. castaneum*, the Y chromosome still persists, containing four small ORF transcripts annotated as ncRNAs, one of which is potentially polycistronic. While the biological roles of these small ORFs remains unclear, future functional studies may determine whether these sequences represent remnants of ancestral large ORFs lost through pseudogenization or if they exemplify a novel mechanism of coding small ORF evolution associated with gene degradation, an evolutionary process that has been recently discussed in the literature [77]. Nonetheless, it is possible that this non-recombining chromosome could, in rare instances, serve as a source for the emergence of new coding fragments, although many may ultimately succumb to decay without selective pressure to maintain their function.

### Deviations from Randomness in Small ORF Lengths Reveal Evolutionary Signatures

The putative small ORF lengths exhibit a significant deviation from the random pattern. Random small ORFs, which results from the random arrangement and rearrangement of nucleotides throughout evolution, exhibit low average sizes in *T. castaneum*, approximately 20 codons (Figure 2B). These patterns are similar to those described for non-conserved and/or potentially non-functional sequences, such as intergenic small ORFs, upstream ORFs, and small ORFs in lncRNAs [39]. In contrast, the putative small ORFs show a more diversified size distribution, as indicated by the median; their average size is higher, approximately 58 codons, a notable departure from the random pattern. Importantly, the vast majority of putative small ORFs are larger than 20 codons (Figure 2B), which suggests that dwarf small ORFs (< 20 codons) are, in fact, scarce.

The average size of conserved putative small ORFs increases as species diverge from *T*. *castaneum*. The 70∼80-codon length observed in small ORFs conserved in distant ancestor groups align with the size described for annotated small ORFs. These findings suggest that larger small ORFs are more likely to become evolutionarily fixed in distantly related taxa.

Regarding small ORF transcript length, previous studies showed that small ORF transcripts are predominantly shorter than canonical mRNAs in metazoans [39]. In *T. castaneum*, putative small ORF transcripts are shorter than the 3.1 kb average size exhibited by canonical mRNAs in metazoans. Notably, at least 214 putative transcripts fall exclusively into alternative small ORF classes, each containing both a small ORF and a larger, potentially canonical CDS.

### Several Small ORFs Can Be Deeply Conserved Across Anciently Rooted Clades

The conservation time offers valuable insights into the adaptive radiations and speciation events to which a small ORF may be related. In this context, we investigated potential orthologs expressed across several groups within the *T. castaneum* evolutionary lineage.

Notably, the existence of these conserved small ORFs across diverse taxa also raises the possibility of horizontal gene transfer (HGT) [78]. In our analysis, we did not detect any prokaryote-conserved small ORFs that were absent in basal eukaryotic lineages and reappeared in more recent ones, a pattern that would otherwise suggest HGT. However, future efforts that compare broader genomic contexts may help clarify whether some of these putative small ORFs disseminated via sporadic transfer events. If confirmed, such scenarios would reinforce the notion that certain small ORFs have deep evolutionary roots, traversing multiple domains of life to fulfill fundamental cellular functions.

From another perspective, the observation that older small ORFs tend to be larger in size suggests that evolutionary constraints may have acted to preserve sequence integrity over vast phylogenetic distances, potentially due to specific structural motifs or functional domains [79,80]. Longer small ORFs might accumulate essential protein folds or binding sites that are more robust to drift, thus enhancing their likelihood of retention in genomes. Thereby, shorter variants are more likely to degenerate rapidly unless they are integrated into critical pathways or acquire specialized functions [81].

In this context, interesting findings were identified in prokaryote-conserved small ORFs. Notably, the six small ORFs conserved in Bacteria point to a potential link to the Last Universal Common Ancestor (LUCA), whose genetic composition and genome organization have been extensively speculated and studied over recent decades [82–86]. A direct link to LUCA would imply that at least some of these small ORFs have undergone stringent purifying selection across billions of years of evolution, preserving crucial functions. Notably, these small ORFs conserved across prokaryotes exceed 50% GC content, suggesting a possible link between GC enrichment and retained functionality. In this context, earlier work in *Drosophila* found that functionally important coding regions often exhibit above-average GC content, likely due to codon usage bias, protein-folding requirements, or selective pressures favoring efficient translation [87]. Thus, it is worth exploring whether these sequences are involved in universal metabolic pathways (e.g., ATP generation and amino acid biosynthesis) or core housekeeping roles (e.g., DNA replication, transcription and translation).

Our functional annotations reveal potential overlaps with such ancient metabolic or housekeeping roles (Additional file 1). Three of the identified Bacteria-conserved small ORFs (XM_015984762.1_6, XM_015979055.1_2, and XM_015978224.1_3) carry gene ontology terms associated with proteolysis, serine-type endopeptidase activity, and acyltransferase activity. Another small ORF (XM_015979099.1_12), which exhibits experimental evidence of translation as assigned by Swiss-Prot, was classified in the pyruvate carboxylase family, with functional implications for pyruvate metabolism and gluconeogenesis (Additional file 1). This connection to a key metabolic enzyme, coupled with translation evidence, underscores the potential relevance of small ORFs to core biochemical pathways.

Furthermore, two of the Bacteria-conserved small ORFs (XM_015984848.1_8 and XM_015979099.1_12) harbor extended protein-binding sites (>10 amino acids), suggesting a notable capacity for stable protein-protein interactions or catalytic complex formation [88] (Additional file 1). The presence of secondary structures comprising β-sheets, α-helices, and disordered regions reinforces the idea that, despite their small size, these peptides may adopt or contribute to functionally significant folds. Notably, the entry XM_015978224.1_3, which also bears translation evidence by ribo-seq, is assigned to gene ontology terms linked to proteolysis. This raises the possibility that small ORFs could engage in fundamental proteolytic pathways conserved since the earliest stages of life.

Consistent with these observations, four additional small ORFs were identified as having a potential common ancestor in Neomura (the clade uniting Archaea and Eukaryotes) (Additional file 1). These four sequences are known small CDSs with experimental evidence of translation. Notably, each small ORF is associated with fundamental cellular machinery: for instance, XM_966256.4_3 and XM_967790.3_2 link to ribosomal protein families (S14 and S27, respectively), while XM_969082.3_2 associates with the Lsm6/SmF family involved in RNA processing, and XM_969699.3_2 relates to the Sec61 translocon complex, an essential component in protein transport [60]. This distribution of core cellular functions echoes the roles proposed for the six bacterial-conserved small ORFs, highlighting that many small ORFs anchor themselves in ancient, crucial pathways, which comprise larger and well-studied gene families.

Detailed structural and mutational analyses, aided by deep protein modeling algorithms, may help determine whether these ancient small ORFs share three-dimensional folds, indicating ancestral protein domains [89]. Such insights could reveal how subtle sequence changes have allowed small ORFs to maintain their adaptive value, whether by preserving core enzymatic activities or by acquiring new binding interfaces relevant to species-specific niches. Interestingly, in our dataset, two Bacteria-conserved small ORFs (XM_015978224.1_3 and XM_015984762.1_6) display predicted peptides with high sequence and tertiary structure similarity across a wide range of taxa (Figure 4B, C, E, F). Both small ORFs appear as isoforms derived from larger genes (Figure 4A, D), a process that can occur through various mechanisms. For instance, in *T. castaneum*, the sequence XM_015978224.1_3 arises via an alternative promoter that yields a transcript containing only two exons, whereas its larger isoform retains four (Figure 4A); importantly, this smaller isoform exhibit evidence of translation assigned by ribo-seq data (Additional file 1). Curiously, these two sequences share conserved regions (Figure 4G) and may act as serine-type paralogs with endopeptidase activity (Additional file 1), assigning their potential to fulfill specialized enzymatic roles despite their shortened structure.

Contrary to early assumptions in the small ORF field [29], these findings demonstrate that several small ORFs exhibit deep conservation, both at the sequence and structural levels. Moreover, these results support the notion that deeply rooted small ORFs are potentially associated with crucial processes, highlighting the need to investigate other highly conserved sequences that remain unannotated but may harbor similarly essential functions. In this context, among the 31 small ORFs originating from the Eukarya ancestor, 13 remain without any assigned protein family or gene ontology annotations, as well as at least nine small ORFs originated in other basal eukaryotic groups. Similarly, two Bacteria-conserved sequences also lack functional annotations (Additional file 1).

### Small ORFs And Gene Duplication Events

Although most duplicated genes become eventually pseudogenes, they can persist, acquire stable functions, and become evolutionarily conserved [90,91]. Furthermore, highly similar gene duplicates offer significant advantages for functional compensation in response to intrinsic inhibitory effects, as well as mutational and environmental challenges [92]. This is especially relevant given that gene duplicates are typically more prone to mutations than their original counterparts [81,93]. Studies in *Drosophila* have linked high levels of duplication and redundancy in developmental genes to phenotypic radiation [90].

In this context, our findings indicate that 70% of the putative small ORFs in *T. castaneum* have potential paralogs in other transcripts (Figure 2F; Additional file 1). Additionally, the pool of potential duplicates with ≥ 80% identity removed during comparative genomic filtering would probably increase this number. Notably, some of these duplications involve large ORFs, which is consistent with the evolutionary model of gene birth after duplication events [77]. According to this model, duplication of canonical coding genes can truncate or rearrange large ORFs, preserving only a small functional core region that remains under selection if it confers a new or modified function. These sequences can lose function and compose pseudogenes or be extended into new large ORFs through a stop codon mutation process, being potentially fixed by natural selection [94,95].

The evolutionary relationship between small ORFs and large ORFs also suggests another convergent process. For example, mechanisms such as alternative transcription, alternative polyadenylation cleavage and alternative splicing can produce new short coding regions that either coexist with the ancestral gene as isoformic small ORFs or evolve independently [10], as exemplified in the figure 4A, D. These isoformic small ORFs effectively mirror the functional domains of their larger counterparts, but may undergo selection for roles that favor more compact or specialized functions [96,97]. A significant portion of 156 entries in our dataset of 454 putative small ORFs are isoformic small ORFs, potentially sharing functional domains with their original counterparts. Curiously, this mirror effect can favor another class of small ORFs, such as the overlapping small ORFs; although not in terms of domain conservation, this effect pertains to sequence conservation, since the evolutionary pressure in one frame can help maintain the codons in other frames, thereby preserving the sequence.

Importantly, gene duplication followed by large CDS fragmentation can result in polycistronic transcripts, where multiple conserved small ORFs can co-occur deriving from these fragments. In *T. castaneum*, the *mlpt* gene exemplifies a well-known case of a polycistronic small ORF arrangement [98]; however, this type of gene is scarce in eukaryotic literature. In contrast, our results reveal that a significant portion of the putative small ORFs co-occur in potential new polycistronic transcripts. Considering that the 35 potential polycistronic transcripts identified encompass 83 (18%) of the putative small ORFs, further studies are required to elucidate if these small ORFs are strictly unfunctional conserved fragments or if some of them are contributing to the birth of new coding genes, following the aforementioned mechanism of gene birth after duplication events.

## CONCLUSION

This study provides compelling evidence that an important amount of small ORFs is far from being evolutionary remnants of “junk DNA” In *T. castaneum.* Our findings demonstrate that a substantial number of these small ORFs are conserved across diverse taxonomic groups, including deeply rooted ancestors such as prokaryotes and the origin of Eukarya.

Importantly, the approaches employed in this study offer a cost-effective strategy for the discovery of small ORFs. By leveraging computational filters and integrating multi-omics data, our methodology provides an accessible and scalable framework that can be combined to other experimental and computational approaches. This integrative strategy facilitates the widespread identification and analysis of small ORFs across several organisms, thereby advancing our understanding of their roles in evolution and development.

Future research should focus on experimental validations of these putative small ORFs to elucidate their precise biological functions and mechanisms. Expanding this integrative approach to other taxa will further enhance our understanding of the evolutionary dynamics and functional significance of small ORFs across the tree of life.

## MATERIAL AND METHODS

All command-line procedures for predicting putative small ORFs in *T. castaneum* were executed locally on Ubuntu operating system (Linux kernel) and are available upon request.

### Intergenic Sequence Retrieving

*De novo* detection of small ORFs was conducted on potentially intergenic sequences of the *T. castaneum* genome [35] (version 5.2) (Bioprojects PRJNA15718 and PRJNA12540), following a similar methodology described in [12]. The multi-FASTA file of the *T. castaneum* genome was retrieved from the NCBI database (National Center for Biotechnology Information, <ncbi.nlm.nih.gov>). Detection was also performed on the complementary strand, obtained by applying the revseq program from the EMBOSS package (version 6.6.0.0) [99] to the *T. castaneum* multi-FASTA genome file. The GFF3 file containing gene location coordinates was also retrieved from the NCBI database; this file was then split into two separate files, containing the coordinates of forward and reverse strand genes, respectively.

The intergenic sequences were retrieved as follows. First, chromosome and scaffold sizes were obtained using the faidx tool from the SAMtools package (version 0.1.19-96b5f2294a) [100]. Second, columns containing chromosome and scaffold identifiers and sizes were extracted from the generated FAI file and saved as a separate file. Third, this file, along with the GFF3 file, was then provided as input to the “complement” tool from the BEDtools package (version v2.26.0) [101], to retrieve intergenic location coordinates. Finally, the intergenic coordinates and the *T. castaneum* genome FASTA file were input into the getfasta tool from BEDtools to generate the intergenic FASTA file.

### *De Novo* Detection of Small ORFs

After retrieving the intergenic FASTA file, *de novo* small ORF detection was performed using the getORF program from the EMBOSS package, configured to extract nucleotide sequences and their corresponding theoretical amino acid chains. The criteria for *de novo* detection were as follows: (a) sequences of 100 codons or fewer, excluding stop codons; (b) sequences of at least seven codons, excluding stop codons; and (c) sequences with in-frame ATG start codons and TAA/TGA/TAG stop codons.

### Comparative Genomic Filter of Small ORFs

The small ORFs identified in the *de novo* detection were subjected to a comparative genomic filter, where potential orthologs were searched in other coleopteran genomes, including *Anoplophora glabripennis*, *Dendroctonus ponderosae*, *Hypothenemus hampei*, and *Leptinotarsa decemlineata*. These coleopterans diverged from *T. castaneum* approximately 162 to 254 million years ago, according to the TimeTree server [102]. The FASTA files of these coleopteran genomes were downloaded from the NCBI database with the following identifiers: GCF_000390285.1 (*A*. *glabripennis*), GCF_000355655.1 (*D*. *ponderosae*), GCA_001012855.1 (*H. hampei*), and GCF_000500325.1 (*L*. *decemlineata*). Subsequently, *de novo* small ORF detection was performed on these genomes using the same parameters previously described, generating a database of small ORFs from other coleopterans. The corresponding amino acid sequences of *T. castaneum* small ORFs were then compared against the amino acid sequences of small ORFs from the other coleopterans using the BLASTp tool from the BLAST+ package (version 2.2.31) [103].

Because the theoretical small ORF peptides are small enough to yield false negative results in BLAST analyses, we modified the search parameters based on strategies applied in [24,98], specifically: (a) the low complexity filter was deactivated; (b) the PAM30 substitution matrix was used; (c) a word size of 2 was applied; and (d) the E-value threshold was set to 0.001.

After BLASTp searches, small ORF-predicted peptides with at least one hit were subjected to a redundancy filter to remove potential duplicates (≥ 80% identity), while retaining the larger sequences. The resulting small ORFs were then compiled into a pool of small ORFs with evidence of evolutionary conservation.

### Refseq Transcript Filter of Small ORFs

The pool of small ORFs with evidence of evolutionary conservation was subjected to BLAST searches against small ORFs extracted from *T. castaneum* RefSeq transcripts using the getORF program. The corresponding amino acid sequences of both small ORF datasets were then submitted to BLASTp searches, using the adjusted parameters for short peptide detection mentioned in the previous section. Putative small ORF-predicted peptides with at least one hit were included in the final pool, which contained small ORFs with evidence of both evolutionary conservation and transcription. This pool is hereafter referred to as *T. castaneum* putative small ORFs. The pool of putative small ORFs was manually annotated with respect to their transcription feature classification.

### Transcriptional Analysis of Potential Orthologs in Eukarya

The putative small ORFs were searched for potential orthologs expressed across the evolutionary scale of *T. castaneum*. First, the full content of RefSeq transcripts available for the multiple taxa within Eukarya were retrieved from the NCBI database using the search commands outlined in Additional file 2. The retrieved file entries are available in Additional file 3.

The phylogeny of Eukarya used as reference for this work was adapted from [104,105]. However, the taxonomic terminology applied in the NCBI searches followed the information provided in the Taxonomy section of the NCBI database, which serves as the reference for NCBI taxonomy.

A local RefSeq transcript database was created for each taxon, containing the available files for their respective species. Subsequently, *de novo* small ORF detection was performed using the getORF program, generating a small ORF database for each taxon. The corresponding amino acid sequences of the putative small ORFs and each taxon database were then compared using BLASTp, configured with the same parameters previously described, but with a less stringent E-value threshold of 0.01. The putative small ORF-predicted peptides with at least one hit were included in the final pool of sequences with evidence of potential orthologs in eukaryotes.

### Transcriptional Analysis of Potential Orthologs in Prokaryotes

Since prokaryotes transcribe most of their genomes and exhibit limited evidence of introns, the detection of transcribed small ORFs in prokaryotes was performed directly on DNA strands. Available multi-FASTA genome files for several prokaryotic species were downloaded from the NCBI RefSeq database using the search commands outlined in Additional file 2. The retrieved file entries are available in Additional file 3. This search resulted in two separate datasets corresponding to Bacteria and Archaea. *De novo* small ORF detection was performed using the getORF program, generating two databases of bacterial and archaeal small ORFs. The corresponding amino acid sequences of the putative small ORFs and the bacterial and archaeal small ORFs were then compared using BLASTp, configured with the same parameters previously described, but with a less stringent E-value threshold of 0.1. Putative small ORF-predicted peptides with at least one hit were included in the pool of sequences with evidence of potential orthologs in prokaryotes.

### Identification of Potential Paralogs Expressed in *T. castaneum*

Two databases of potential paralogs were constructed, one for small ORFs and another for large ORFs (> 100 codons). The small ORF database was composed of small ORFs extracted from RefSeq transcripts, that were discarded in the transcriptional filter step. The large ORFs were obtained by applying the getORF program to the *T. castaneum* RefSeq transcripts. The corresponding amino acid sequences of putative small ORFs were then compared against the amino acid sequences of ORFs in both databases using BLASTp, configured with the same parameters previously described and an E-value threshold of 0.001. The predicted amino acid sequences of small and large ORFs with at least one hit were included in the pool of potential paralogs expressed in *T. castaneum*.

### Identification of Potential Polycistronic Small ORF Transcripts

Potential polycistronic small ORF transcripts were identified based on the repeated occurrence of small ORF transcript identifiers.

### Transcript Length Analysis

The lengths of small ORF transcripts, along with the mean, median, standard deviation, maximum, and minimum values, were calculated using the faSize program from Jim Kent’s Utilities [106]

### Small ORF Length Analysis

The lengths of small ORFs, including the mean, median, standard deviation, maximum, and minimum values, were determined using the faSize program [106].

### GC Content, Codon and Amino Acid Usage

The codon and amino acid usage, as well as the average GC content per codon and mean GC values for each nucleotide position within codons, were analyzed using the Cusp tool from the EMBOSS package. These analyses were conducted on the nucleotide sequences of putative small ORFs, random small ORFs and genomic CDSs of *T. castaneum*, retrieved from the Assembly section of the NCBI database.

### Functional Enrichment

Functional analysis of small ORF-predicted peptides was performed using the InterPro server, which classified them into families, predicted their domains and important sites, and assigned them Gene Ontology (GO) terms [107].

### Expression patterns in *T. castaneum*

Fourteen publicly available RNA-seq libraries of *T. castaneum* were retrieved from the NCBI Sequence Read Archive (SRA). Libraries were selected based on the following criteria: (i) coverage of the widest possible range of developmental stages, (ii) absence of biotic or abiotic treatments, and (iii) exclusion of datasets involving gene silencing experiments. The selected datasets comprised: five embryonic libraries (SRR3310884, SRR3310889, SRR3310897, SRR3310901, and SRR3310903), two libraries from adult females (SRR28121203 and SRR28205973), two ovarian libraries (SRR19548811 and SRR19548810), one male library (SRR28205972), two testis libraries (SRR19548808 and SRR19548809), and two larval libraries (SRR21991470 and SRR21991471). Raw sequence data were converted to FASTQ format using the SRA Toolkit (v3.1.1) [108]. Adapter trimming and quality filtering were performed using Trimmomatic (v0.39) [109], with quality filtering based on the Phred33 score and parameters tailored to each sequencing technology. Transcript quantification was conducted using Salmon (v1.10.1) [110]. An index was created with the salmon index function using the reference transcriptome of *T. castaneum* (Tcas5.2, GCF_000002335.3) and the small ORF transcripts identified in this study. Expression levels were then estimated as transcripts per million (TPM) via the salmon quant function. TPM values were recovered for each gene across all libraries. Finally, the expression data were visualized using Circos plots [111] to illustrate the transcriptional landscape of small ORFs across developmental stages and tissues.

Fourteen public RNAseq libraries of *T. castaneum* were recovered from NCBI/SRA (https://www.ncbi.nlm.nih.gov/sra). The selection criteria of the libraries were: libraries representing the greatest number of developmental stages possible of *T. castaneum*, libraries without coming from experiments influenced by treatments whether biotic or abiotic and libraries without coming from experiments with gene silencing.

### Translation Evidence Analysis

Translation evidence analysis was conducted by submitting the corresponding amino acid sequences of *T. castaneum* putative small ORFs to BLASTp searches against amino acid sequences of small ORFs detected by ribo-seq, available in the sORF.org database [112], as well as sequences in the Swiss-Prot database [113], using the same modified parameters used in the comparative and transcriptional filters.

### Structural Analysis of Small ORF-Predicted Peptides

The tertiary structure of small ORF-predicted peptides showed in figure 5 were predicted using the Falcon2 web server [114]. The global and local stereochemical quality of all predicted models were performed by Rampage [115], PROCHECK [116], ProSA [117], VoroMQA [118], ProQ3D [119], Qprob [120], and SVMQA [121]. The best structural models were refined by ModRefiner [122]. The global alignment of small ORF-predicted peptides in figure 5 were performed with Clustal [123].

**Figure 5.**
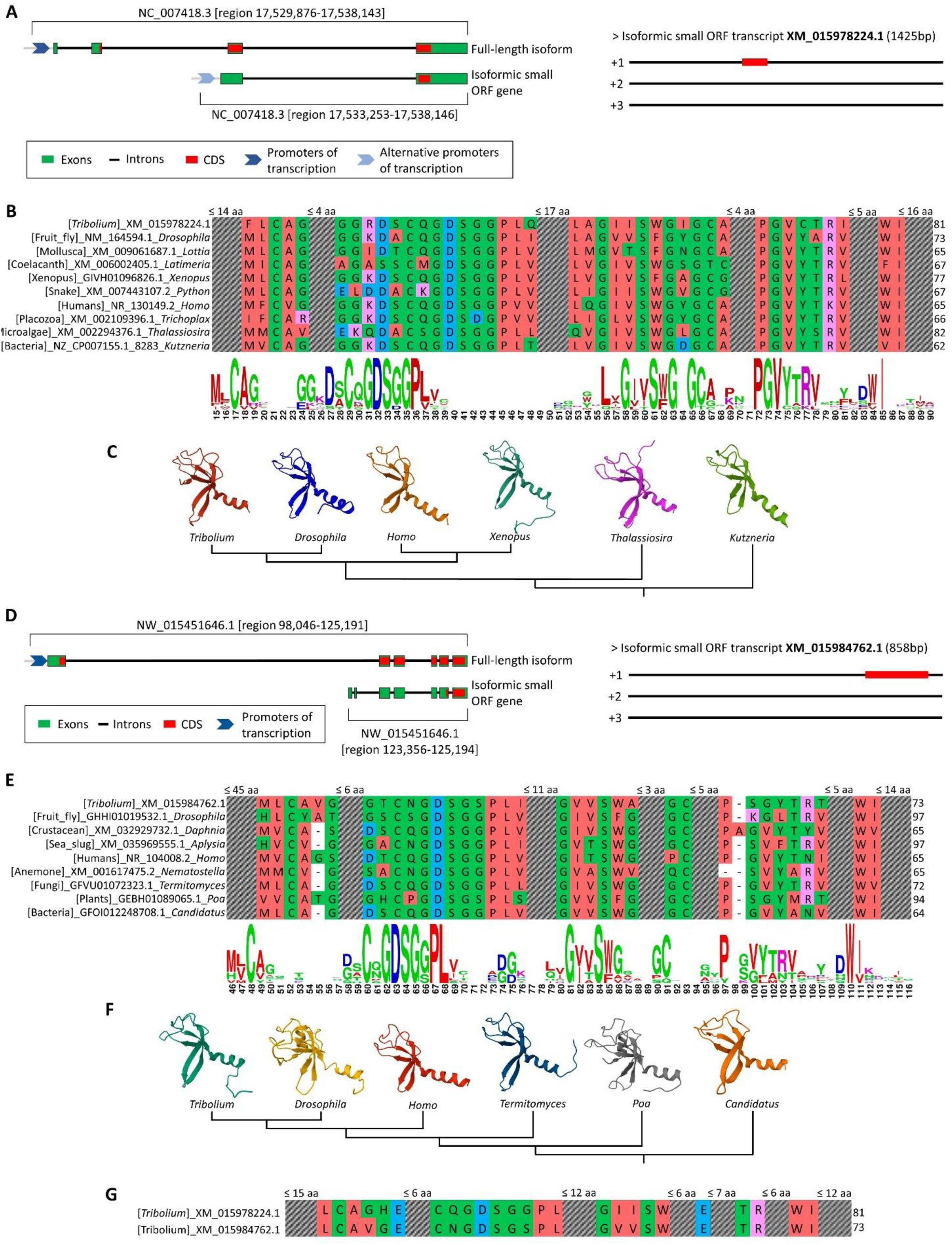
Conservation analysis of two bacteria-conserved small ORF-predicted peptides in *T. castaneum*. **(A)** Genomic context and gene structure of the first small ORF in *T. castaneum* (left), alongside its transcript localization (right). **(B)** Global alignment of the first peptide with putative orthologs across diverse taxa, followed by its sequence logo. **(C)** Phylogenetic tree showing the evolutionary relationships of the first peptide, alongside its predicted tertiary structure. **(D)** Genomic context and gene structure of the second small ORF in *T. castaneum* (left), alongside its transcript localization (right). **(E)** Global alignment of the second peptide with putative orthologs, accompanied by its sequence logo. **(F)** Phylogenetic tree of the second peptide across taxa, shown with its predicted tertiary structure. **(G)** Pairwise alignment of both bacteria-conserved peptides, highlighting sequence similarity and suggesting a potential paralogous relationship.

### Negative Control

The negative control was conducted by submitting random small ORF to the prediction filters used in this study. These random small ORFs were generated by applying the getORF program to the reverse and non-complementary sequence of the *T. castaneum* genome. The reverse genome sequence was obtained using the revseq program on the *T. castaneum* multi-FASTA genome file. The BLASTp filtering steps were repeated multiple times with the same parameters as previously described, but with variable E-values of 0.001, 0.01, 0.1, 1, and 10.

### Ethics approval and consent to participate

Not applicable.

### Consent for publication

Not applicable.

### Availability of data and materials

All data generated or analyzed during this study are included in this published article (and its supplementary information files) or available from the corresponding author on reasonable request.

## Authors’ contributions

DG-A conceived the study, performed most analyses, and wrote the manuscript. RN-d-F supervised the project and guided both analyses and manuscript preparation. DAT contributed to the methodological design. EP-C performed structural prediction analyses. JJM-M conducted the expression analyses using RNA-seq libraries. MFN contributed to manuscript writing and critical revision.

## Acknowledgements

We gratefully acknowledge Eva Guerra and Guido Guerra for their valuable support throughout the development of this research

