## Additional file 2 for "The small ORFeome of *Tribolium castaneum* reveals deeply conserved small ORFs broadly expressed across tissues"

**Table 1**. Search commands used to retrieve the available RefSeq transcript files for each taxon on the *Tribolium castaneum* phylogenetic scale of life. Each retrieved file corresponds to the RefSeq transcripts of a unique species.

| Taxon | Search commands | Retrieved files |
|---|---|---|
| Coleoptera | "Coleoptera"[Organism] AND ("latest refseq"[filter] AND all[filter] NOT anomalous[filter]) | 6 |
| Insecta | "Insecta"[Organism] NOT "Coleoptera"[Organism] AND "latest refseq"[filter] | 33 |
| Hexapoda | "Hexapoda"[Organism] NOT "Insecta"[Organism] AND "latest refseq"[filter] | 1 |
| Pancrustacea | "Pancrustacea"[Organism] NOT "Hexapoda"[Organism] AND "latest refseq"[filter] | 2 |
| Arthropoda | "Arthropoda"[Organism] NOT "Pancrustacea"[Organism] AND "latest refseq"[filter] | 8 |
| Ecdysozoa | "Ecdysozoa"[Organism] NOT "Arthropoda"[Organism] AND "latest refseq"[filter] | 7 |
| Protostomia | "Protostomia"[Organism] NOT "Ecdysozoa"[Organism] AND "latest refseq"[filter] | 14 |
| Eubilateria | "Deuterostomia"[Organism] AND "latest refseq"[filter] | 129 |
| Neuralia | "Cnidaria"[Organism] AND "latest refseq"[filter] | 6 |
| Eumetazoa | "Eumetazoa"[Organism] NOT "Cnidaria"[Organism] NOT "Bilateria"[Organism] | 1 |
| Metazoa | "Metazoa"[Organism] NOT "Eumetazoa"[Organism] AND (latest[filter] AND all[filter] NOT anomalous[filter]) | 1 |
| Opisthokonta | "Opisthokonta"[Organism] NOT "Metazoa"[Organism] AND (latest[filter] AND all[filter] NOT anomalous[filter]) | 70 |
| Amorphea | "Amoebozoa"[Organism] AND (latest[filter] AND all[filter] NOT anomalous[filter]) | 11 |
| Eukarya | "Eukaryota"[Organism] NOT "Amoebozoa"[Organism] NOT "Opisthokonta"[Organism] AND (latest[filter] AND all[filter] NOT anomalous[filter]) | 148 |
| | **Total** | **437** |

**Table 2**. Search commands used to retrieve the available RefSeq transcript files for Archaea and Bacteria. Each retrieved file corresponds to the RefSeq genome of a unique species.

| Taxon | Search commands | Retrieved files |
|---|---|---|
| Archaea | ("Archaea"[Organism] OR Archaea[All Fields]) AND ("latest refseq"[filter] AND all[filter] NOT anomalous[filter]) | 740 |

| | | |
|---|---|---|
| Bacteria | ("Bacteria"[Organism] OR Bacteria[All Fields]) AND ("latest refseq"[filter] AND "complete genome"[filter] AND "representative genome"[filter] AND all[filter] NOT anomalous[filter]) | 1.545 |
| | Total | 2.285 |

**Table 3** – GC content per codon in *Tribolium castaneum*.

| | GC content | | | Average percentage |
|---|---|---|---|---|
| | First base | Second base | Third base | |
| Putative small ORFs | 45.13% | 42.88% | 46.58% | 44.87% |
| Random small ORFs | 40.68% | 43.52% | 46.95% | 43.72% |
| *Tribolium* genomic CDSs | 49.52% | 38.14% | 51.09% | 46.25% |

**Table 4** – GC content per codon of putative small ORFs in *Tribolium castaneum*, categorized by their conservation levels.

| | GC content | | | Average percentage |
|---|---|---|---|---|
| | First base | Second base | Third base | |
| *Tribolium* | 45.13% | 42.88% | 46.58% | 44.87% |
| Coleoptera | 46.07% | 46.02% | 49.20% | 47.10% |
| Insecta | 46.65% | 44.63% | 50.27% | 47.18% |
| Hexapoda | 47.87% | 42.91% | 53.78% | 48.19% |
| Pancrustcea | 48.43% | 43.62% | 54.42% | 48.82% |
| Arthropoda | 46.45% | 40.90% | 51.81% | 46.39% |
| Ecdysozoa | 47.54% | 41.28% | 52.49% | 47.10% |
| Protostomia | 48.36% | 43.87% | 50.93% | 47.72% |
| Eubilateria | 46.91% | 42.68% | 49.99% | 46.53% |
| Neuralia | 46.32% | 42.20% | 49.64% | 46.05% |
| Eumetazoa | 47.14% | 43.30% | 54.27% | 48.24% |

| | | | | |
|---|---|---|---|---|
| Metazoa | 47.90% | 39.12% | 52.96% | 46.66% |
| Opisthokonta | 49.57% | 41.92% | 52.31% | 47.93% |
| Amorphea | 48.59% | 43.70% | 53.38% | 48.56% |
| Eukarya | 48.19% | 42.03% | 50.17% | 46.80% |
| Neomura | 49.90% | 48.13% | 56.97% | 51.67% |
| Neomura + Bacteria | 55.17% | 45.81% | 50.29% | 50.42% |

**Table 5.** Amino acid usage of predicted peptide/protein sequences of ORFs in *Tribolium castaneum*.

| Amino acids | Putative small ORFs | Randon small ORFs | Large ORFs |
|---|---|---|---|
| A | 54.43 | 42.76 | 58.73 |
| V | 57.63 | 54.56 | 63.69 |
| L | 87.65 | 97.68 | 91.29 |
| I | 58.72 | 58.91 | 56.10 |
| P | 52.56 | 50.48 | 54.92 |
| M | 37.59 | 60.49 | 20.81 |
| F | 47.92 | 50.86 | 38.85 |
| W | 18.09 | 19.54 | 10.83 |
| G | 53.61 | 44.88 | 53.78 |
| S | 81.02 | 86.61 | 78.52 |
| T | 61.61 | 67.44 | 60.02 |
| C | 31.19 | 33.36 | 19.59 |
| N | 45.58 | 43.21 | 49.72 |
| Q | 36.15 | 35.75 | 43.09 |
| Y | 29.79 | 26.45 | 30.56 |
| D | 32.60 | 20.15 | 53.08 |
| E | 44.61 | 27.13 | 69.56 |
| K | 65.47 | 64.77 | 69.36 |
| R | 76.31 | 90.16 | 53.32 |
| H | 27.49 | 24.81 | 24.19 |

**Table 6**. Codon relative frequency of the putative small ORFs of *Tribolium castaneum* categorized by conservation levels.

| | | Putative small ORFs of *Tribolium castaneum* | | | | | | | | | | | | | | | |
|---|---|---|---|---|---|---|---|---|---|---|---|---|---|---|---|---|---|
| Codon | Amino acid | Coleoptera | Insecta | Hexapoda | Pancrustacea | Arthropoda | Ecdysozoa | Protostomia | Eubilateria | Neuralia | Eumetazoa | Metazoa | Opisthokonta | Amorphea | Eukarya | Archaea | Bacteria |
| GCA | A | 15.93 | 14.64 | 16.95 | 12.99 | 11.57 | 8.13 | 12.41 | 13.04 | 12.31 | 5.48 | 10.34 | 6.27 | 9.43 | 11.14 | 11.79 | 11.70 |
| GCC | A | 14.60 | 16.67 | 23.15 | 21.54 | 20.43 | 22.21 | 20.14 | 18.45 | 21.60 | 17.23 | 19.53 | 19.70 | 23.99 | 19.15 | 33.40 | 29.24 |
| GCG | A | 12.49 | 11.97 | 12.40 | 11.40 | 9.53 | 11.92 | 11.51 | 10.50 | 11.56 | 13.31 | 9.77 | 12.99 | 12.00 | 10.79 | 11.79 | 5.85 |
| GCT | A | 14.25 | 14.09 | 14.06 | 16.16 | 15.15 | 17.34 | 17.08 | 16.38 | 16.08 | 14.10 | 14.36 | 21.05 | 16.28 | 17.76 | 23.58 | 21.44 |
| TGC | C | 16.70 | 16.85 | 16.95 | 19.96 | 17.70 | 21.67 | 17.98 | 16.70 | 17.84 | 24.28 | 17.81 | 20.15 | 28.28 | 18.45 | 19.65 | 9.75 |
| TGT | C | 17.05 | 17.22 | 13.64 | 15.52 | 17.02 | 18.96 | 17.98 | 17.17 | 20.35 | 22.71 | 14.93 | 17.47 | 22.28 | 19.15 | 11.79 | 17.54 |
| GAC | D | 15.16 | 16.94 | 19.84 | 19.64 | 22.13 | 18.96 | 18.34 | 16.38 | 16.33 | 17.23 | 20.10 | 18.81 | 18.00 | 18.45 | 7.86 | 15.60 |
| GAT | D | 17.05 | 17.59 | 18.60 | 20.91 | 20.09 | 20.31 | 19.96 | 18.76 | 20.35 | 24.28 | 23.55 | 23.74 | 23.14 | 21.59 | 23.58 | 19.49 |
| GAA | E | 26.10 | 28.82 | 29.35 | 26.61 | 30.13 | 32.23 | 30.03 | 31.17 | 29.89 | 24.28 | 32.17 | 35.38 | 31.71 | 34.47 | 21.61 | 35.09 |
| GAG | E | 14.53 | 17.41 | 19.43 | 21.86 | 19.40 | 19.50 | 18.34 | 18.13 | 17.84 | 17.23 | 26.42 | 19.70 | 16.28 | 17.76 | 11.79 | 15.60 |
| TTC | F | 18.74 | 18.23 | 21.50 | 21.22 | 22.13 | 25.46 | 22.65 | 21.47 | 22.61 | 23.49 | 19.53 | 19.70 | 20.57 | 20.54 | 23.58 | 23.39 |
| TTT | F | 24.28 | 23.48 | 19.02 | 21.22 | 23.66 | 21.13 | 21.94 | 24.17 | 23.36 | 18.79 | 22.98 | 19.70 | 14.57 | 23.33 | 11.79 | 9.75 |
| GGA | G | 17.89 | 19.25 | 14.06 | 22.17 | 19.57 | 22.48 | 19.78 | 20.83 | 21.35 | 23.49 | 17.81 | 19.26 | 21.42 | 20.89 | 23.58 | 38.99 |
| GGC | G | 13.47 | 14.37 | 17.78 | 19.05 | 15.15 | 15.71 | 16.90 | 13.68 | 16.33 | 22.71 | 12.64 | 13.88 | 22.28 | 17.76 | 29.47 | 21.44 |
| GGG | G | 13.96 | 15.38 | 15.71 | 17.11 | 15.15 | 15.44 | 17.44 | 16.38 | 17.08 | 22.71 | 16.08 | 14.78 | 18.00 | 14.28 | 13.75 | 9.75 |
| GGT | G | 12.14 | 12.71 | 10.34 | 11.40 | 12.77 | 14.90 | 13.66 | 12.88 | 13.31 | 10.18 | 15.51 | 12.09 | 15.42 | 11.49 | 11.79 | 21.44 |
| CAC | H | 14.03 | 14.37 | 13.64 | 14.25 | 16.17 | 17.88 | 15.10 | 15.11 | 15.32 | 19.58 | 13.79 | 20.60 | 18.00 | 18.45 | 19.65 | 23.39 |
| CAT | H | 12.98 | 13.91 | 10.75 | 9.50 | 12.09 | 13.27 | 15.46 | 13.99 | 12.06 | 14.10 | 12.64 | 14.78 | 14.57 | 16.02 | 9.82 | 13.65 |
| ATA | I | 15.65 | 13.54 | 11.99 | 11.40 | 13.45 | 11.92 | 13.31 | 13.68 | 12.81 | 18.01 | 10.91 | 10.30 | 12.00 | 13.93 | 7.86 | 11.70 |
| ATC | I | 15.65 | 16.48 | 20.67 | 21.22 | 19.92 | 21.13 | 17.62 | 18.92 | 18.84 | 15.66 | 18.96 | 18.36 | 19.71 | 16.71 | 31.43 | 27.29 |
| ATT | I | 22.95 | 23.85 | 20.26 | 22.49 | 23.15 | 24.11 | 24.81 | 26.08 | 24.87 | 21.14 | 21.25 | 19.70 | 21.42 | 20.89 | 13.75 | 11.70 |
| AAA | K | 39.79 | 39.23 | 44.23 | 38.64 | 48.51 | 46.32 | 44.95 | 44.36 | 49.99 | 42.29 | 53.42 | 45.68 | 49.70 | 51.18 | 37.33 | 50.68 |
| AAG | K | 23.79 | 25.79 | 26.87 | 27.56 | 24.68 | 22.48 | 23.55 | 25.28 | 21.35 | 25.84 | 26.42 | 24.63 | 22.28 | 21.24 | 21.61 | 1.95 |
| CTA | L | 9.40 | 9.21 | 9.10 | 7.29 | 8.85 | 7.58 | 9.53 | 9.54 | 7.79 | 6.27 | 9.77 | 10.30 | 6.00 | 10.79 | 5.89 | 3.90 |
| CTC | L | 12.21 | 12.06 | 13.64 | 12.67 | 11.40 | 11.65 | 11.69 | 12.09 | 11.30 | 12.53 | 10.91 | 12.54 | 9.43 | 12.88 | 31.43 | 29.24 |
| CTG | L | 14.17 | 15.29 | 17.78 | 16.47 | 12.43 | 13.27 | 13.31 | 12.88 | 13.06 | 16.45 | 14.36 | 13.88 | 13.71 | 11.14 | 9.82 | 7.80 |
| CTT | L | 12.98 | 12.43 | 14.88 | 11.09 | 10.72 | 12.46 | 12.41 | 11.61 | 13.56 | 10.18 | 9.19 | 11.64 | 9.43 | 9.75 | 3.93 | 13.65 |
| TTA | L | 15.65 | 16.12 | 12.82 | 12.35 | 14.47 | 13.27 | 12.23 | 14.95 | 15.32 | 11.75 | 15.51 | 12.99 | 7.71 | 18.11 | 7.86 | 11.70 |
| TTG | L | 20.98 | 22.65 | 23.56 | 23.44 | 24.17 | 25.46 | 22.11 | 23.06 | 24.62 | 19.58 | 27.00 | 26.42 | 23.14 | 20.20 | 13.75 | 7.80 |
| ATG | M | 35.93 | 35.55 | 31.42 | 32.63 | 35.57 | 36.02 | 33.44 | 34.98 | 34.92 | 37.59 | 34.46 | 33.59 | 36.85 | 33.77 | 29.47 | 23.39 |

| Codon | Amino acid | Coleoptera | Insecta | Hexapoda | Pancrustacea | Arthropoda | Ecdysozoa | Protostomia | Eubilateria | Neuralia | Eumetazoa | Metazoa | Opisthokonta | Amorphea | Eukarya | Archaea | Bacteria |
|---|---|---|---|---|---|---|---|---|---|---|---|---|---|---|---|---|---|
| AAC | N | 20.21 | 20.81 | 21.91 | 24.71 | 23.49 | 23.84 | 19.60 | 21.78 | 20.35 | 22.71 | 24.70 | 24.63 | 25.71 | 22.63 | 21.61 | 23.39 |
| AAT | N | 20.84 | 21.55 | 18.19 | 18.37 | 21.79 | 19.50 | 17.08 | 19.56 | 23.61 | 19.58 | 22.40 | 17.02 | 19.71 | 19.50 | 13.75 | 9.75 |
| CCA | P | 16.98 | 14.55 | 14.06 | 16.16 | 11.75 | 9.21 | 14.20 | 14.79 | 11.30 | 10.18 | 15.51 | 13.44 | 12.85 | 12.19 | 7.86 | 9.75 |
| CCC | P | 14.25 | 16.12 | 15.71 | 18.69 | 17.87 | 15.71 | 16.72 | 15.42 | 17.33 | 21.93 | 16.08 | 17.02 | 22.28 | 16.71 | 21.61 | 19.49 |
| CCG | P | 14.81 | 14.18 | 14.88 | 15.52 | 11.40 | 11.11 | 13.13 | 12.72 | 9.29 | 11.75 | 10.34 | 12.99 | 9.43 | 13.93 | 11.79 | 13.65 |
| CCT | P | 12.00 | 12.43 | 10.34 | 12.35 | 10.55 | 10.83 | 12.41 | 11.93 | 11.30 | 12.53 | 8.04 | 12.54 | 14.57 | 11.84 | 13.75 | 17.54 |
| CAA | Q | 21.40 | 20.35 | 23.15 | 20.91 | 20.26 | 21.67 | 20.32 | 23.06 | 19.84 | 21.93 | 21.83 | 20.15 | 15.42 | 22.63 | 27.51 | 27.29 |
| CAG | Q | 13.61 | 13.63 | 11.99 | 15.84 | 14.13 | 13.54 | 14.38 | 13.20 | 12.56 | 16.45 | 22.40 | 17.91 | 16.28 | 15.32 | 11.79 | 15.60 |
| AGA | R | 18.32 | 16.48 | 12.82 | 14.57 | 14.64 | 14.63 | 16.18 | 16.22 | 14.07 | 19.58 | 14.93 | 14.33 | 17.14 | 18.80 | 19.65 | 15.60 |
| AGG | R | 15.44 | 15.75 | 18.60 | 16.16 | 15.15 | 15.44 | 15.46 | 14.79 | 15.32 | 16.45 | 14.93 | 16.57 | 18.00 | 15.32 | 23.58 | 13.65 |
| CGA | R | 16.56 | 14.55 | 14.47 | 11.72 | 10.89 | 12.19 | 13.84 | 12.09 | 12.56 | 13.31 | 9.77 | 13.88 | 15.42 | 9.75 | 3.93 | 0.00 |
| CGC | R | 10.25 | 8.84 | 8.68 | 9.82 | 9.53 | 8.94 | 9.35 | 8.59 | 7.79 | 8.61 | 9.77 | 11.20 | 6.00 | 9.75 | 13.75 | 11.70 |
| CGG | R | 11.79 | 10.04 | 7.03 | 8.24 | 8.34 | 7.86 | 10.25 | 8.75 | 8.04 | 7.83 | 6.32 | 9.40 | 6.86 | 8.36 | 5.89 | 9.75 |
| CGT | R | 11.16 | 9.95 | 7.44 | 5.07 | 6.64 | 8.40 | 8.63 | 9.54 | 6.53 | 3.13 | 5.17 | 9.40 | 7.71 | 9.05 | 13.75 | 15.60 |
| AGC | S | 12.42 | 11.51 | 11.99 | 11.09 | 11.23 | 10.29 | 9.35 | 10.65 | 9.80 | 9.40 | 8.04 | 11.64 | 5.14 | 11.14 | 11.79 | 13.65 |
| AGT | S | 14.25 | 12.16 | 14.06 | 14.25 | 11.92 | 12.19 | 11.87 | 12.88 | 13.06 | 8.61 | 10.91 | 9.85 | 6.86 | 8.36 | 13.75 | 19.49 |
| TCA | S | 16.63 | 14.92 | 12.40 | 10.45 | 12.77 | 10.56 | 13.13 | 13.20 | 12.31 | 7.05 | 6.89 | 11.64 | 6.86 | 11.84 | 11.79 | 13.65 |
| TCC | S | 13.47 | 12.80 | 13.64 | 12.35 | 11.06 | 11.38 | 12.77 | 11.93 | 11.81 | 17.23 | 15.51 | 9.85 | 10.28 | 11.84 | 25.54 | 19.49 |
| TCG | S | 15.44 | 15.47 | 13.64 | 14.57 | 13.11 | 14.09 | 14.56 | 15.11 | 12.31 | 13.31 | 13.79 | 15.67 | 12.85 | 16.37 | 17.68 | 9.75 |
| TCT | S | 12.07 | 11.79 | 11.16 | 8.55 | 8.68 | 7.31 | 9.35 | 10.97 | 9.80 | 10.18 | 9.19 | 11.20 | 11.14 | 9.40 | 17.68 | 3.90 |
| ACA | T | 19.86 | 18.14 | 15.71 | 14.57 | 15.15 | 14.36 | 16.36 | 15.11 | 17.33 | 13.31 | 13.21 | 13.44 | 12.00 | 13.93 | 15.72 | 15.60 |
| ACC | T | 16.07 | 16.30 | 15.71 | 13.62 | 15.15 | 12.73 | 14.38 | 14.95 | 12.56 | 11.75 | 12.64 | 11.64 | 12.85 | 13.58 | 17.68 | 11.70 |
| ACG | T | 17.05 | 14.83 | 15.30 | 13.62 | 11.92 | 11.65 | 12.59 | 13.20 | 11.30 | 10.18 | 17.23 | 10.75 | 12.00 | 9.75 | 9.82 | 7.80 |
| ACT | T | 13.40 | 13.17 | 10.34 | 10.77 | 10.72 | 11.11 | 11.87 | 12.40 | 13.06 | 10.96 | 8.04 | 11.64 | 12.00 | 11.84 | 5.89 | 17.54 |
| GTA | V | 9.82 | 10.04 | 8.68 | 7.29 | 9.19 | 9.48 | 9.17 | 8.75 | 10.30 | 9.40 | 12.06 | 8.96 | 10.28 | 11.14 | 13.75 | 7.80 |
| GTC | V | 12.84 | 13.45 | 15.30 | 16.47 | 16.85 | 18.15 | 15.28 | 16.54 | 16.08 | 14.10 | 14.36 | 17.02 | 14.57 | 17.06 | 21.61 | 33.14 |
| GTG | V | 14.67 | 14.18 | 19.84 | 18.06 | 15.49 | 15.71 | 16.54 | 15.11 | 16.58 | 15.66 | 20.68 | 14.78 | 20.57 | 14.62 | 13.75 | 11.70 |
| GTT | V | 17.19 | 17.13 | 15.71 | 16.16 | 18.89 | 17.34 | 16.36 | 16.86 | 16.58 | 13.31 | 17.81 | 15.67 | 14.57 | 14.97 | 13.75 | 21.44 |
| TGG | W | 19.58 | 19.16 | 16.12 | 16.79 | 16.51 | 14.09 | 17.44 | 15.58 | 17.33 | 19.58 | 16.08 | 13.44 | 15.42 | 15.67 | 7.86 | 11.70 |
| TAC | Y | 13.75 | 15.66 | 23.15 | 18.69 | 20.94 | 21.67 | 17.44 | 17.65 | 17.33 | 20.36 | 18.96 | 18.81 | 23.14 | 18.11 | 25.54 | 25.34 |
| TAT | Y | 13.40 | 14.00 | 13.64 | 14.89 | 16.85 | 11.92 | 14.38 | 14.15 | 18.59 | 17.23 | 10.34 | 13.44 | 14.57 | 12.54 | 11.79 | 9.75 |
