## Additional file 3 for "The small ORFeome of *Tribolium castaneum* reveals deeply conserved small ORFs broadly expressed across tissues"

**ADDITIONAL FILE 3** – NCBI RefSeq transcriptome and genome entries used in the comparative analyses of potential orthologs.

**Coleoptera**
GCF_000355655.1
GCF_000390285.2
GCF_000500325.1
GCF_000648695.1
GCF_001412225.1
GCF_001937115.1

**Insecta**
GCF_000001215.4
GCF_000002195.4
GCF_000002325.3
GCF_000005115.1
GCF_000005135.1
GCF_000005155.2
GCF_000005175.2
GCF_000005575.2
GCF_000005975.2
GCF_000006295.1
GCF_000142985.2
GCF_000143395.1
GCF_000151625.1
GCF_000184785.2
GCF_000187915.1
GCF_000188095.2
GCF_000204515.1
GCF_000214255.1
GCF_000217595.1
GCF_000220905.1
GCF_000269505.1
GCF_000330985.1
GCF_000341935.1
GCF_000475195.1
GCF_000696155.1
GCF_000754195.2
GCF_000836215.1
GCF_001277935.1
GCF_001654015.1
GCF_001654025.1
GCF_002204515.2
GCF_002706865.1
GCF_002891405.2

**Hexapoda**
GCF_002217175.1

**Pancrustacea**
GCF_000591075.1
GCF_000764305.1

**Arthropoda**
GCF_000208615.1
GCF_000239435.1
GCF_000255335.1
GCF_000365465.2
GCF_000517525.1
GCF_000671375.1
GCF_002443255.1
GCF_002532875.1

**Ecdysozoa**
GCF_000002985.6
GCF_000149515.1
GCF_000181795.1
GCF_000183805.2
GCF_000485595.1
GCF_000507365.1
GCF_001040885.1
GCF_000242695.1
GCF_002880755.1

**Protostomia**
GCF_000002075.1
GCF_000237925.1
GCF_000297895.1
GCF_000326865.1
GCF_000327385.1
GCF_000457365.1
GCF_000524195.1
GCF_000699445.1
GCF_000715545.1
GCF_001039355.2
GCF_001194135.1
GCF_002022765.2
GCF_002113885.1
GCF_003073045.1

**Eubilateria**
GCF_000001405.38
GCF_000001635.26
GCF_000001895.5
GCF_000001905.1
GCF_000002035.6
GCF_000002235.4
GCF_000002275.2
GCF_000002285.3
GCF_000002295.2
GCF_000002315.5
GCF_000003025.6
GCF_000003605.2
GCF_000003625.3
GCF_000003815.1
GCF_000004195.3
GCF_000004335.2
GCF_000004665.1
GCF_000090745.1
GCF_000146605.2
GCF_000146795.2
GCF_000147115.1
GCF_000151735.1
GCF_000151805.1
GCF_000151845.1
GCF_000151885.1
GCF_000151905.2
GCF_000164805.1
GCF_000164845.2
GCF_000165045.1
GCF_000165445.2
GCF_000180615.1
GCF_000181275.1
GCF_000181295.1
GCF_000181335.3
GCF_000186305.1
GCF_000189315.1
GCF_000208655.1
GCF_000215625.1
GCF_000223135.1
GCF_000224145.3
GCF_000225785.1
GCF_000230535.1
GCF_000233375.1
GCF_000235385.1
GCF_000236235.1
GCF_000238935.1
GCF_000238955.4
GCF_000239375.1
GCF_000239395.1
GCF_000239415.1
GCF_000241765.3
GCF_002872995.1
GCF_900095145.1
GCF_000243295.1
GCF_000247695.1
GCF_000247795.1
GCF_000247815.1
GCF_000258655.2
GCF_000260255.1
GCF_000260355.1
GCF_000264685.3
GCF_000276665.1
GCF_000277835.1
GCF_000280705.1
GCF_000283155.1
GCF_000292845.1
GCF_000296735.1
GCF_000296755.1
GCF_000298275.1
GCF_000298355.1
GCF_000298735.2
GCF_000299155.1
GCF_000308155.1
GCF_000311805.1
GCF_000313985.1
GCF_000317375.1
GCF_000321225.1
GCF_000325575.1
GCF_000327345.1
GCF_000331425.1
GCF_000331955.2
GCF_000334495.1
GCF_000337955.1
GCF_000337975.1
GCF_000344595.1
GCF_000349665.1
GCF_000349705.1
GCF_000355885.1
GCF_000364345.1
GCF_000372685.2
GCF_000385455.1
GCF_000400835.1
GCF_000409795.2
GCF_000412655.1
GCF_000442215.1
GCF_000455745.1
GCF_000464555.1
GCF_000485575.1
GCF_000493695.1
GCF_000500345.1
GCF_000523025.1
GCF_000534875.1
GCF_000633615.1
GCF_000721915.3
GCF_000772875.2
GCF_000951615.1
GCF_001465895.1
GCF_001522545.2
GCF_001577835.1
GCF_001660625.1
GCF_001663975.1
GCF_001704415.1
GCF_001858045.2
GCF_002021735.1
GCF_002078875.1
GCF_002163495.1
GCF_002201575.1
GCF_002234675.1
GCF_002263795.1
GCF_002775205.1
GCF_002863925.1
GCF_000226395.1
GCF_000715135.1
GCF_002880775.1
GCF_002901205.1
GCF_002910315.2
GCF_003121395.1
GCF_003255815.1
GCF_900094665.1

**Neuralia**
GCF_002571385.1
GCF_000004095.1
GCF_000209225.1
GCF_000222465.1
GCF_001417965.1
GCF_002042975.1

**Eumetazoa**
GCF_000150275.1

**Metazoa**
GCF_000090795.1

**Opisthokonta**
GCF_000001985.1
GCF_900079805.1
GCF_000002495.2
GCF_000002515.2
GCF_000002525.2
GCF_000002545.3
GCF_000002655.1
GCF_000002715.2
GCF_000002855.3
GCF_000002865.3
GCF_000002945.1
GCF_000003125.1
GCF_000003515.1
GCF_000003835.1
GCF_000003855.2
GCF_000004155.1
GCF_000006275.2
GCF_000006335.3
GCF_000006445.2
GCF_000026365.1
GCF_000026945.1
GCF_000027005.1
GCF_000091025.4
GCF_000091045.1
GCF_000142805.1
GCF_000143105.1
GCF_000143185.1
GCF_000143365.1
GCF_000143535.2
GCF_000146045.2
GCF_000146465.1
GCF_000149205.2
GCF_000149555.1
GCF_000149955.1
GCF_000150675.1
GCF_000151355.1
GCF_000182925.2
GCF_000182965.3
GCF_000184455.2
GCF_000185945.1
GCF_000187245.1
GCF_000188695.1
GCF_000209165.1
GCF_000219625.1
GCF_000226095.1
GCF_000226115.1
GCF_000151665.1
GCF_000710875.1
GCF_000226545.1
GCF_000227115.2
GCF_000235365.1
GCF_000236905.1
GCF_000237345.1
GCF_000240135.3
GCF_000243375.1
GCF_000277815.2
GCF_000280035.1
GCF_000303195.2
GCF_000304475.1
GCF_000315875.1
GCF_000315915.1
GCF_000328475.2
GCF_000348985.1
GCF_000388065.1
GCF_001298625.1
GCF_001417885.1
GCF_001640025.1
GCF_001653235.2
GCF_001672515.1
GCF_002742065.1
GCF_900007375.1

**Amorphea**
GCF_000787575.1
GCF_000004695.1
GCF_000004825.1
GCF_000142905.1
GCF_000190715.1
GCF_000203815.1
GCF_000208925.1
GCF_000209125.1
GCF_000257125.1
GCF_000313135.1
GCF_000330505.1

**Eukarya**
GCF_900002335.2
GCF_000001735.4
GCF_000002415.2
GCF_000002425.4
GCF_000002455.1
GCF_000002765.4
GCF_000002775.4
GCF_000002975.1
GCF_000003195.3
GCF_000003225.3
GCF_000003745.3
GCF_000004075.2
GCF_000004255.2
GCF_000005005.2
GCF_000005505.3
GCF_000006355.1
GCF_000006405.1
GCF_000006515.1
GCF_000006565.2
GCF_000018645.1
GCF_000091205.1
GCF_000141845.1
GCF_000142945.1
GCF_000143415.4
GCF_000148765.1
GCF_000149405.2
GCF_000149755.1
GCF_000150535.2
GCF_000150955.2
GCF_000151545.1
GCF_900002375.1
GCF_900002385.1
GCF_000151685.1
GCF_000165345.1
GCF_000165365.1
GCF_000165395.1
GCF_000165425.1
GCF_000184155.1
GCF_000186865.1
GCF_000189635.1
GCF_000194455.1
GCF_000208745.1
GCF_000208865.1
GCF_000219495.3
GCF_000220395.1
GCF_000223845.1
GCF_000226075.1
GCF_000231095.1
GCF_000240725.1
GCF_000247585.1
GCF_000258005.1
GCF_000263155.2
GCF_000281045.1
GCF_000313045.1
GCF_000313855.2
GCF_000315295.1
GCF_000315625.1
GCF_000317415.1
GCF_000321355.1
GCF_000327365.1
GCF_000331145.1
GCF_000340665.1
GCF_000341285.1
GCF_000342415.1
GCF_000346465.2
GCF_000346735.1
GCF_000350225.1
GCF_000365185.1
GCF_000372725.1
GCF_000375325.1
GCF_000390325.2
GCF_000393655.1
GCF_000413155.1
GCF_000414095.1
GCF_000442705.1
GCF_000463585.1
GCF_000471905.2
GCF_000478725.1
GCF_000493195.1
GCF_000495115.1
GCF_000499385.1
GCF_000499425.1
GCF_000499545.2
GCF_000499605.1
GCF_000499745.1
GCF_000499845.1
GCF_000504015.1
GCF_000511025.2
GCF_000512975.1
GCF_000520075.1
GCF_000520115.1
GCF_000524495.1
GCF_000612285.1
GCF_000612305.1
GCF_000633955.1
GCF_000686985.2
GCF_000691945.2
GCF_000695525.1
GCF_000696525.1
GCF_000709005.1

GCF_000740895.1
GCF_000741045.1
GCF_000743755.1
GCF_000769155.1
GCF_000801105.1
GCF_000816755.2
GCF_000817695.2
GCF_000826755.1
GCF_000956335.1
GCF_000981445.1
GCF_000987745.1
GCF_001190045.1
GCF_001263595.1
GCF_001406875.1
GCF_001411555.1
GCF_001433935.1
GCF_001531365.1
GCF_001540865.1
GCF_001601855.1
GCF_001602025.1
GCF_001625215.1
GCF_001654055.1
GCF_001659605.1
GCF_001680005.1
GCF_001683475.1
GCF_001865875.1
GCF_001876935.1
GCF_001879085.1
GCF_001879475.1
GCF_001957025.1
GCF_001995035.1
GCF_002007265.1
GCF_002127325.1
GCF_002168275.1
GCF_002207925.1
GCF_002211085.1
GCF_002303985.1
GCF_002738345.1
GCF_002738365.1
GCF_002742605.1
GCF_002806865.1
GCF_002870075.1
GCF_002906115.1
GCF_002994745.1
GCF_900000015.1

**Archaea**

GCF_900215215.1
GCF_900215575.1
GCF_900289035.1
GCF_000006175.1
GCF_000006805.1
GCF_000007005.1
GCF_000007065.1
GCF_000007305.1
GCF_000007345.1
GCF_000008265.1
GCF_000008645.1
GCF_000008665.1
GCF_000009185.1
GCF_000009965.1
GCF_000011085.1
GCF_000011105.1
GCF_000011185.1
GCF_000011205.1
GCF_000011585.1
GCF_000012285.1
GCF_000012545.1
GCF_000013445.1
GCF_000013725.1
GCF_000015145.1
GCF_000015205.1
GCF_000015225.1
GCF_000015765.1
GCF_000015825.1
GCF_000015945.1
GCF_000016125.1
GCF_000016525.1
GCF_000016605.1
GCF_000017165.1
GCF_000017185.1
GCF_000017225.1
GCF_000017625.1
GCF_000017945.1
GCF_000018305.1
GCF_000018365.1
GCF_000018485.1
GCF_000019605.1
GCF_000020905.1
GCF_000021965.1
GCF_000022205.1
GCF_000022365.1
GCF_000022385.1
GCF_000022405.1
GCF_000022425.1
GCF_000022465.1
GCF_000022485.1
GCF_000022545.1
GCF_000023945.1
GCF_000023965.1
GCF_000023985.1
GCF_000024185.1
GCF_000024305.1
GCF_000024625.1
GCF_000024745.1
GCF_000025285.1
GCF_000025325.1
GCF_000025505.1
GCF_000025525.1
GCF_000025625.1
GCF_000025665.1
GCF_000025685.1
GCF_000025865.1
GCF_000026045.1
GCF_000063445.1
GCF_000069025.1
GCF_000091665.1
GCF_000092185.1
GCF_000092305.1
GCF_000092465.1
GCF_000144915.1
GCF_000145295.1
GCF_000147875.1
GCF_000148385.1
GCF_000151085.1
GCF_000151105.2
GCF_000151205.2
GCF_000151225.1
GCF_000151245.1
GCF_000152265.2
GCF_000166095.1
GCF_000172995.2
GCF_000175555.1
GCF_000179575.2
GCF_000186365.1
GCF_000187225.1
GCF_000189555.1
GCF_000189795.1
GCF_000189815.1
GCF_000189835.1
GCF_000189855.1
GCF_000189895.1
GCF_000189915.1
GCF_000189935.1
GCF_000189955.1
GCF_000189975.1
GCF_000189995.1
GCF_000190015.1
GCF_000190035.1
GCF_000190055.1
GCF_000190075.1
GCF_000190095.1
GCF_000190115.1
GCF_000190135.1
GCF_000190155.1
GCF_000190175.1
GCF_000190315.1
GCF_000191585.1
GCF_000193375.1
GCF_000194625.1
GCF_000195895.1
GCF_000195915.1
GCF_000195935.2
GCF_000196655.1
GCF_000196895.1
GCF_000204415.1
GCF_000204925.1
GCF_000211475.1
GCF_000213215.1
GCF_000214415.1
GCF_000214725.1
GCF_000215915.2
GCF_000215995.1
GCF_000217715.1
GCF_000217995.1
GCF_000220175.1
GCF_000220645.1
GCF_000221185.1
GCF_000223395.1
GCF_000223905.1
GCF_000224475.1
GCF_000226975.2
GCF_000230715.2
GCF_000230735.2
GCF_000230955.2
GCF_000231015.2
GCF_000235565.1
GCF_000235685.2
GCF_000237865.1
GCF_000241145.1
GCF_000242875.2
GCF_000243255.1
GCF_000243315.1
GCF_000243455.1
GCF_000245095.1
GCF_000245115.1
GCF_000245135.1
GCF_000245155.1
GCF_000245175.1
GCF_000245195.1
GCF_000245215.1
GCF_000245235.1
GCF_000245255.1
GCF_000245275.1
GCF_000246985.2
GCF_000251105.1
GCF_000253055.1
GCF_000258515.1
GCF_000259215.1
GCF_000263735.1
GCF_000264495.1
GCF_000265525.1
GCF_000275605.1
GCF_000275865.1
GCF_000281695.1
GCF_000283335.1
GCF_000296615.1
GCF_000299395.1
GCF_000300255.2
GCF_000302455.1
GCF_000304355.2
GCF_000306725.1
GCF_000306765.2
GCF_000308215.1
GCF_000317795.1
GCF_000320505.1
GCF_000327485.1
GCF_000327505.1
GCF_000328285.1
GCF_000328665.1
GCF_000328685.1
GCF_000328925.1
GCF_000328945.1
GCF_000334895.1
GCF_000336615.1
GCF_000336635.1
GCF_000336655.1
GCF_000336675.1
GCF_000336695.1
GCF_000336715.1
GCF_000336735.1
GCF_000336755.1
GCF_000336775.1
GCF_000336795.1
GCF_000336815.1
GCF_000336835.1
GCF_000336855.1
GCF_000336875.1
GCF_000336895.1
GCF_000336915.1
GCF_000336935.1
GCF_000336955.1
GCF_000336975.1
GCF_000336995.1
GCF_000337015.1
GCF_000337035.1
GCF_000337055.1
GCF_000337075.1
GCF_000337095.1
GCF_000337115.1
GCF_000337135.1
GCF_000337155.1
GCF_000337175.1
GCF_000337195.1
GCF_000337215.1
GCF_000337235.1
GCF_000337255.1
GCF_000337275.1
GCF_000337295.1
GCF_000337315.1
GCF_000337335.1
GCF_000337355.1
GCF_000337375.1
GCF_000337395.1
GCF_000337415.1
GCF_000337435.1
GCF_000337455.1
GCF_000337475.1
GCF_000337495.1
GCF_000337515.1
GCF_000337535.1
GCF_000337555.1
GCF_000337575.1
GCF_000337595.1
GCF_000337615.1
GCF_000337635.1
GCF_000337655.1
GCF_000337675.1
GCF_000337695.1
GCF_000337715.1
GCF_000337735.1
GCF_000337755.1
GCF_000337775.1
GCF_000337795.1
GCF_000337815.1
GCF_000337835.1
GCF_000337855.1
GCF_000337875.1
GCF_000337895.1
GCF_000337915.1
GCF_000338775.1
GCF_000340315.1
GCF_000350305.1
GCF_000363885.1
GCF_000364745.1
GCF_000371805.1
GCF_000376445.1
GCF_000376965.1
GCF_000379085.1
GCF_000383975.1
GCF_000385565.1
GCF_000400975.1
GCF_000403645.1
GCF_000404165.1
GCF_000404225.1
GCF_000421805.1
GCF_000427685.1
GCF_000430485.1
GCF_000430905.1
GCF_000446015.1
GCF_000447865.2
GCF_000455345.1
GCF_000455365.1
GCF_000470655.1
GCF_000474235.1
GCF_000485535.1
GCF_000493245.1
GCF_000495475.1
GCF_000499765.1
GCF_000504205.1
GCF_000504565.1
GCF_000513315.1
GCF_000513435.1
GCF_000513855.1
GCF_000517445.1
GCF_000517625.1
GCF_000529525.1
GCF_000585495.1
GCF_000591055.1
GCF_000621965.1
GCF_000632495.1
GCF_000685155.1
GCF_000685395.1
GCF_000685635.1
GCF_000690595.1
GCF_000691505.1
GCF_000691865.1
GCF_000698785.1
GCF_000700025.1
GCF_000710605.1
GCF_000710615.1
GCF_000711215.1
GCF_000711905.1
GCF_000723185.1
GCF_000723845.1
GCF_000725425.1
GCF_000730175.1
GCF_000731985.1
GCF_000734035.1
GCF_000739065.1
GCF_000739555.1
GCF_000739575.1
GCF_000739595.1
GCF_000744315.1
GCF_000744455.1
GCF_000745485.1
GCF_000746075.1
GCF_000746205.1
GCF_000747605.1
GCF_000755225.1
GCF_000755245.1
GCF_000762265.1
GCF_000765475.1
GCF_000769655.1
GCF_000784335.1
GCF_000784355.1
GCF_000789255.1
GCF_000800805.1
GCF_000812185.1
GCF_000813245.1
GCF_000816105.1
GCF_000824705.1
GCF_000827835.1
GCF_000828575.1
GCF_000949015.1
GCF_000953115.1
GCF_000955905.1
GCF_000956175.1
GCF_000966265.1
GCF_000968355.1
GCF_000968395.1
GCF_000968435.1
GCF_000969885.1
GCF_000969905.1
GCF_000969945.1
GCF_000969965.1
GCF_000969985.1
GCF_000970005.1
GCF_000970025.1
GCF_000970045.1
GCF_000970065.1
GCF_000970085.1
GCF_000970145.1
GCF_000970165.1
GCF_000970185.1
GCF_000970205.1
GCF_000970225.1
GCF_000970245.1
GCF_000970265.1
GCF_000970285.1
GCF_000970305.1
GCF_000970325.1
GCF_000978935.1
GCF_000978945.1
GCF_000978955.1
GCF_000978965.1
GCF_000979015.1
GCF_000979025.1
GCF_000979035.1
GCF_000979045.1

GCF_000979095.1
GCF_000979105.1
GCF_000979115.1
GCF_000979125.1
GCF_000979175.1
GCF_000979185.1
GCF_000979195.1
GCF_000979205.1
GCF_000979255.1
GCF_000979265.1
GCF_000979275.1
GCF_000979295.1
GCF_000979335.1
GCF_000979345.1
GCF_000979375.1
GCF_000979385.1
GCF_000979415.1
GCF_000979425.1
GCF_000979455.1
GCF_000979475.1
GCF_000979495.1
GCF_000979505.1
GCF_000979515.1
GCF_000979555.1
GCF_000979575.1
GCF_000979585.1
GCF_000979595.1
GCF_000979635.1
GCF_000979655.1
GCF_000979665.1
GCF_000979675.1
GCF_000979685.1
GCF_000979735.1
GCF_000979745.1
GCF_000979755.1
GCF_000979765.1
GCF_000979815.1
GCF_000979825.1
GCF_000979845.1
GCF_000979855.1
GCF_000979895.1
GCF_000979915.1
GCF_000979925.1
GCF_000979935.1
GCF_000979975.1
GCF_000979995.1
GCF_000980005.1
GCF_000980025.1
GCF_000980055.1
GCF_000980075.1
GCF_000980085.1
GCF_000980105.1
GCF_000980135.1
GCF_000980155.1
GCF_000980175.1
GCF_000993805.1
GCF_001006045.1
GCF_001006085.1
GCF_001011115.1
GCF_001017125.1
GCF_001027005.1
GCF_001190965.1
GCF_001261915.1
GCF_001266655.1
GCF_001266675.1
GCF_001266695.1
GCF_001266715.1
GCF_001266735.1
GCF_001280425.1
GCF_001280455.1
GCF_001282785.1
GCF_001304615.1
GCF_001305655.1
GCF_001307315.1
GCF_001315885.1
GCF_001315945.1
GCF_001316045.1
GCF_001316065.1
GCF_001316085.1
GCF_001317345.1
GCF_001368915.1
GCF_001399695.1
GCF_001402935.1
GCF_001402945.1
GCF_001412615.1
GCF_001418715.1
GCF_001433455.1
GCF_001458655.1
GCF_001462205.1
GCF_001462395.1
GCF_001469865.1
GCF_001469875.2
GCF_001469955.1
GCF_001477655.1
GCF_001481295.1
GCF_001481635.1
GCF_001481685.1
GCF_001482285.1
GCF_001483125.1
GCF_001484195.1
GCF_001484685.1
GCF_001485535.1
GCF_001485555.1
GCF_001485575.1
GCF_001488575.1
GCF_001541925.1
GCF_001542905.1
GCF_001548675.1
GCF_001559955.1
GCF_001559965.1
GCF_001559975.1
GCF_001559985.1
GCF_001560035.1
GCF_001560045.1
GCF_001560065.1
GCF_001560085.1
GCF_001560115.1
GCF_001560125.1
GCF_001560135.1
GCF_001560165.1
GCF_001560195.1
GCF_001560205.1
GCF_001560215.1
GCF_001560245.1
GCF_001560275.1
GCF_001560285.1
GCF_001560305.1
GCF_001560315.1
GCF_001560355.1
GCF_001560375.1
GCF_001560385.1
GCF_001560405.1
GCF_001560435.1
GCF_001560455.1
GCF_001560465.1
GCF_001560485.1
GCF_001560515.1
GCF_001560525.1
GCF_001560555.1
GCF_001560565.1
GCF_001560595.1
GCF_001560605.1
GCF_001560635.1
GCF_001560645.1
GCF_001560675.1
GCF_001560685.1
GCF_001560715.1
GCF_001560725.1
GCF_001560755.1
GCF_001560765.1
GCF_001560795.1
GCF_001560815.1
GCF_001560835.1
GCF_001560915.1
GCF_001563245.1
GCF_001571385.1
GCF_001571405.1
GCF_001577775.1
GCF_001592435.1
GCF_001593955.1
GCF_001602375.1
GCF_001625445.1
GCF_001639265.1
GCF_001639275.1
GCF_001639285.1
GCF_001639295.1
GCF_001647085.1
GCF_001647155.1
GCF_001663375.1
GCF_001719125.1
GCF_001723155.1
GCF_001729285.1
GCF_001729375.1
GCF_001729385.1
GCF_001729395.1
GCF_001729455.1
GCF_001729965.1
GCF_001748385.1
GCF_001767315.1
GCF_001861355.1
GCF_001886955.1
GCF_001888095.1
GCF_001889405.1
GCF_001950595.1
GCF_001953745.1
GCF_001971705.1
GCF_001989615.1
GCF_002025255.1
GCF_002072215.1
GCF_002077075.2
GCF_002078205.2
GCF_002078355.1
GCF_002114285.1
GCF_002135005.1
GCF_002135045.1
GCF_002153915.1
GCF_002156705.1
GCF_002156965.1
GCF_002177135.1
GCF_002194565.1
GCF_002197185.1
GCF_002201915.1
GCF_002208625.1
GCF_002214165.1
GCF_002214365.1
GCF_002214385.1
GCF_002214465.1
GCF_002214485.1
GCF_002214505.1
GCF_002214525.1
GCF_002214545.1
GCF_002214565.1
GCF_002214585.1
GCF_002214605.1
GCF_002215285.1
GCF_002215305.1
GCF_002215405.1
GCF_002215445.1
GCF_002215485.1
GCF_002215525.1
GCF_002215565.1
GCF_002243045.1
GCF_002252585.1
GCF_002252725.1
GCF_002252735.1
GCF_002252745.1
GCF_002252755.1
GCF_002252805.1
GCF_002252815.1
GCF_002252835.1
GCF_002252865.1
GCF_002252875.1
GCF_002252895.1
GCF_002252925.1
GCF_002252965.1
GCF_002252985.1
GCF_002286985.1
GCF_002287175.1
GCF_002287195.1
GCF_002287215.1
GCF_002287235.1
GCF_002355635.1
GCF_002355655.1
GCF_002379115.1
GCF_002487355.1
GCF_002494345.1
GCF_002572525.1
GCF_002727095.1
GCF_002727125.1
GCF_002761295.1
GCF_002787055.1
GCF_002788215.1
GCF_002813085.1
GCF_002813655.1
GCF_002813675.1
GCF_002813695.1
GCF_002844195.1
GCF_002844335.1
GCF_002855455.1
GCF_002906215.1
GCF_002906575.1
GCF_002945325.1
GCF_002973515.1
GCF_003045125.1
GCF_003058365.1
GCF_003116855.1
GCF_003149675.1
GCF_003173335.1
GCF_003173355.1
GCF_003175215.1
GCF_003201675.1
GCF_003201765.1
GCF_003201835.1
GCF_003205235.1
GCF_003264935.1
GCF_003265405.1
GCF_003268005.1
GCF_003269155.1
GCF_003287355.1
GCF_003298465.1
GCF_003336245.1
GCF_003342675.1
GCF_003342695.1
GCF_003344565.1
GCF_003351065.1
GCF_003369835.1
GCF_003382685.1
GCF_003382985.1
GCF_003383005.1
GCF_003383015.1
GCF_003383025.1
GCF_003383065.1
GCF_003383085.1
GCF_003383095.1
GCF_003383195.1
GCF_003385755.1
GCF_003400025.1
GCF_003400145.1
GCF_900003505.1
GCF_900003515.1
GCF_900003525.1
GCF_900003565.1
GCF_900003575.1
GCF_900004275.1
GCF_900005775.1
GCF_900012635.1
GCF_900036045.1
GCF_900064395.1
GCF_900069765.1
GCF_900079115.1
GCF_900079125.1
GCF_900090055.1
GCF_900095295.1
GCF_900095385.1
GCF_900095815.1
GCF_900100335.1
GCF_900100385.1
GCF_900100715.1
GCF_900100875.1
GCF_900102305.1
GCF_900103505.1
GCF_900103715.1
GCF_900104065.1
GCF_900106715.1
GCF_900106905.1
GCF_900107195.1
GCF_900107205.1
GCF_900107665.1
GCF_900108095.1
GCF_900108165.1
GCF_900108505.1
GCF_900109065.1
GCF_900109425.1
GCF_900109595.1
GCF_900109695.1
GCF_900110215.1
GCF_900110455.1
GCF_900110465.1
GCF_900110535.1
GCF_900110865.1
GCF_900111485.1
GCF_900111645.1
GCF_900111935.1
GCF_900112175.1
GCF_900112205.1
GCF_900113245.1
GCF_900114025.1
GCF_900114435.1
GCF_900114455.1
GCF_900114585.1
GCF_900114835.1
GCF_900115675.1
GCF_900115785.1
GCF_900116205.1
GCF_900129775.1
GCF_900142335.1
GCF_900143675.1
GCF_900156425.1
GCF_900156445.1
GCF_900156475.1
GCF_900167955.1
GCF_900176435.1
GCF_900177455.1
GCF_900188065.1
GCF_900188075.1
GCF_900196725.1
GCF_900198835.1

**Bacteria**

GCF_900170005.1
GCF_900184705.1
GCF_900183405.1
GCF_000005825.2
GCF_000006605.1
GCF_000006685.1
GCF_000006725.1
GCF_000007025.1
GCF_000007085.1
GCF_000007125.1
GCF_000007365.1
GCF_000007485.1
GCF_000007605.1
GCF_000007625.1
GCF_000007705.1
GCF_000007725.1
GCF_000007745.1
GCF_000007865.1
GCF_000007905.1
GCF_000007945.1
GCF_000008025.1
GCF_000008045.1
GCF_000008205.1
GCF_000008325.1
GCF_000008345.1
GCF_000008365.1
GCF_000008385.1
GCF_000008465.1
GCF_000008885.1
GCF_000009125.1
GCF_000009145.1
GCF_000009365.1
GCF_000009545.1
GCF_000009765.2
GCF_000009825.1
GCF_000009845.1
GCF_000009865.1
GCF_000009905.1
GCF_000009945.1
GCF_000009985.1
GCF_000010065.1
GCF_000010125.1
GCF_000010145.1
GCF_000010165.1
GCF_000010185.1
GCF_000010285.1
GCF_000010305.1
GCF_000010405.1
GCF_000010425.1
GCF_000010505.1
GCF_000010525.1

| | | | | | |
|---|---|---|---|---|---|
| GCF_000010605.1 | GCF_000014425.1 | GCF_000018405.1 | GCF_000022565.1 | GCF_000025705.1 | GCF_000143985.1 |
| GCF_000010625.1 | GCF_000014445.1 | GCF_000018605.1 | GCF_000022725.1 | GCF_000025725.1 | GCF_000144605.1 |
| GCF_000010665.1 | GCF_000014465.1 | GCF_000018665.1 | GCF_000022905.1 | GCF_000025845.1 | GCF_000144625.1 |
| GCF_000010785.1 | GCF_000014505.1 | GCF_000018685.1 | GCF_000022965.1 | GCF_000025885.1 | GCF_000144645.1 |
| GCF_000010985.1 | GCF_000014565.1 | GCF_000018785.1 | GCF_000023025.1 | GCF_000025905.1 | GCF_000144675.1 |
| GCF_000011025.1 | GCF_000014705.1 | GCF_000018885.1 | GCF_000023065.1 | GCF_000025925.1 | GCF_000144695.1 |
| GCF_000011225.1 | GCF_000014725.1 | GCF_000018945.1 | GCF_000023105.1 | GCF_000025945.1 | GCF_000145035.1 |
| GCF_000011245.1 | GCF_000014745.1 | GCF_000019165.1 | GCF_000023125.1 | GCF_000025965.1 | GCF_000145235.1 |
| GCF_000011305.1 | GCF_000014765.1 | GCF_000019205.1 | GCF_000023145.1 | GCF_000026005.1 | GCF_000145255.1 |
| GCF_000011465.1 | GCF_000014785.1 | GCF_000019225.1 | GCF_000023205.1 | GCF_000026125.1 | GCF_000145275.1 |
| GCF_000011485.1 | GCF_000014825.1 | GCF_000019285.1 | GCF_000023225.1 | GCF_000026405.1 | GCF_000145615.1 |
| GCF_000011645.1 | GCF_000014865.1 | GCF_000019345.1 | GCF_000023245.1 | GCF_000026505.1 | GCF_000146025.2 |
| GCF_000011685.1 | GCF_000014885.1 | GCF_000019405.1 | GCF_000023265.1 | GCF_000026605.1 | GCF_000146185.1 |
| GCF_000011905.1 | GCF_000014925.1 | GCF_000019585.2 | GCF_000023285.1 | GCF_000027225.1 | GCF_000146505.1 |
| GCF_000011965.2 | GCF_000014965.1 | GCF_000019665.1 | GCF_000023325.1 | GCF_000027325.1 | GCF_000147355.1 |
| GCF_000012085.2 | GCF_000015025.1 | GCF_000019685.1 | GCF_000023445.1 | GCF_000046685.1 | GCF_000147695.2 |
| GCF_000012145.1 | GCF_000015045.1 | GCF_000019725.1 | GCF_000023465.1 | GCF_000046705.1 | GCF_000147715.2 |
| GCF_000012225.1 | GCF_000015125.1 | GCF_000019745.1 | GCF_000023605.1 | GCF_000055785.1 | GCF_000147835.2 |
| GCF_000012305.1 | GCF_000015245.1 | GCF_000019785.1 | GCF_000023705.1 | GCF_000055945.1 | GCF_000148645.1 |
| GCF_000012325.1 | GCF_000015285.1 | GCF_000019845.1 | GCF_000023745.1 | GCF_000056065.1 | GCF_000152825.2 |
| GCF_000012385.1 | GCF_000015305.1 | GCF_000019905.1 | GCF_000023785.1 | GCF_000058485.1 | GCF_000153485.2 |
| GCF_000012405.1 | GCF_000015345.1 | GCF_000019945.1 | GCF_000023825.1 | GCF_000060345.1 | GCF_000154785.2 |
| GCF_000012425.1 | GCF_000015445.1 | GCF_000019965.1 | GCF_000023845.1 | GCF_000062885.1 | GCF_000155735.2 |
| GCF_000012465.1 | GCF_000015505.1 | GCF_000020005.1 | GCF_000023865.1 | GCF_000063545.1 | GCF_000157355.2 |
| GCF_000012485.1 | GCF_000015565.1 | GCF_000020025.1 | GCF_000023885.1 | GCF_000063605.1 | GCF_000157895.3 |
| GCF_000012565.1 | GCF_000015585.1 | GCF_000020045.1 | GCF_000023905.1 | GCF_000067165.1 | GCF_000158275.2 |
| GCF_000012585.1 | GCF_000015645.1 | GCF_000020065.1 | GCF_000023925.1 | GCF_000067205.1 | GCF_000163895.2 |
| GCF_000012605.1 | GCF_000015665.1 | GCF_000020145.1 | GCF_000024005.1 | GCF_000069225.1 | GCF_000164675.2 |
| GCF_000012645.1 | GCF_000015725.1 | GCF_000020165.1 | GCF_000024025.1 | GCF_000069945.1 | GCF_000164695.2 |
| GCF_000012665.1 | GCF_000015745.1 | GCF_000020205.1 | GCF_000024065.1 | GCF_000069965.1 | GCF_000164865.1 |
| GCF_000012685.1 | GCF_000015865.1 | GCF_000020225.1 | GCF_000024085.1 | GCF_000070465.1 | GCF_000164985.3 |
| GCF_000012725.1 | GCF_000016065.1 | GCF_000020305.1 | GCF_000024105.1 | GCF_000072485.1 | GCF_000165465.1 |
| GCF_000012745.1 | GCF_000016085.1 | GCF_000020365.1 | GCF_000024125.1 | GCF_000085865.1 | GCF_000165485.1 |
| GCF_000012765.1 | GCF_000016165.1 | GCF_000020385.1 | GCF_000024205.1 | GCF_000090965.1 | GCF_000165505.1 |
| GCF_000012805.1 | GCF_000016185.1 | GCF_000020465.1 | GCF_000024225.1 | GCF_000091125.1 | GCF_000165715.2 |
| GCF_000012825.1 | GCF_000016285.1 | GCF_000020485.1 | GCF_000024265.1 | GCF_000091305.1 | GCF_000166055.1 |
| GCF_000012845.1 | GCF_000016345.1 | GCF_000020505.1 | GCF_000024285.1 | GCF_000091325.1 | GCF_000166135.1 |
| GCF_000012865.1 | GCF_000016405.1 | GCF_000020525.1 | GCF_000024325.1 | GCF_000091465.1 | GCF_000166395.1 |
| GCF_000012885.1 | GCF_000016425.1 | GCF_000020545.1 | GCF_000024345.1 | GCF_000091565.1 | GCF_000166415.1 |
| GCF_000012925.1 | GCF_000016505.1 | GCF_000020605.1 | GCF_000024365.1 | GCF_000091785.1 | GCF_000172635.2 |
| GCF_000012965.1 | GCF_000016545.1 | GCF_000020625.1 | GCF_000024385.1 | GCF_000091985.1 | GCF_000175215.2 |
| GCF_000012985.1 | GCF_000016565.1 | GCF_000020645.1 | GCF_000024405.1 | GCF_000092045.1 | GCF_000175255.2 |
| GCF_000013005.1 | GCF_000016645.1 | GCF_000020685.1 | GCF_000024425.1 | GCF_000092105.1 | GCF_000175575.2 |
| GCF_000013025.1 | GCF_000016745.1 | GCF_000020725.1 | GCF_000024505.1 | GCF_000092125.1 | GCF_000176855.2 |
| GCF_000013125.1 | GCF_000016765.1 | GCF_000020945.1 | GCF_000024545.1 | GCF_000092205.1 | GCF_000176915.2 |
| GCF_000013145.1 | GCF_000016825.1 | GCF_000020965.1 | GCF_000024565.1 | GCF_000092225.1 | GCF_000177235.2 |
| GCF_000013165.1 | GCF_000016905.1 | GCF_000021025.1 | GCF_000024605.1 | GCF_000092245.1 | GCF_000177535.2 |
| GCF_000013285.1 | GCF_000016985.1 | GCF_000021285.1 | GCF_000024725.1 | GCF_000092265.1 | GCF_000177615.2 |
| GCF_000013325.1 | GCF_000017005.1 | GCF_000021325.1 | GCF_000024765.1 | GCF_000092285.1 | GCF_000177635.2 |
| GCF_000013345.1 | GCF_000017245.1 | GCF_000021345.1 | GCF_000024785.1 | GCF_000092365.1 | GCF_000178115.2 |
| GCF_000013365.1 | GCF_000017265.1 | GCF_000021385.1 | GCF_000024845.1 | GCF_000092385.1 | GCF_000178835.2 |
| GCF_000013405.1 | GCF_000017305.1 | GCF_000021485.1 | GCF_000024865.1 | GCF_000092405.1 | GCF_000178875.2 |
| GCF_000013605.1 | GCF_000017405.1 | GCF_000021545.1 | GCF_000024885.1 | GCF_000092425.1 | GCF_000178955.2 |
| GCF_000013645.1 | GCF_000017425.1 | GCF_000021565.1 | GCF_000024925.1 | GCF_000092505.1 | GCF_000178975.2 |
| GCF_000013665.1 | GCF_000017465.2 | GCF_000021685.1 | GCF_000024945.1 | GCF_000092585.1 | GCF_000179035.2 |
| GCF_000013745.1 | GCF_000017545.1 | GCF_000021725.1 | GCF_000024965.1 | GCF_000092645.1 | GCF_000179395.2 |
| GCF_000013765.1 | GCF_000017565.1 | GCF_000021745.1 | GCF_000024985.1 | GCF_000092785.1 | GCF_000179635.2 |
| GCF_000013885.1 | GCF_000017585.1 | GCF_000021865.1 | GCF_000025005.1 | GCF_000092825.1 | GCF_000179915.2 |
| GCF_000013945.1 | GCF_000017645.1 | GCF_000021885.1 | GCF_000025065.1 | GCF_000092845.1 | GCF_000180175.2 |
| GCF_000013985.1 | GCF_000017685.1 | GCF_000021925.1 | GCF_000025085.1 | GCF_000092865.1 | GCF_000182745.2 |
| GCF_000014025.1 | GCF_000017705.1 | GCF_000021945.1 | GCF_000025185.1 | GCF_000092905.1 | GCF_000183135.1 |
| GCF_000014045.1 | GCF_000017805.1 | GCF_000021985.1 | GCF_000025205.1 | GCF_000092925.1 | GCF_000183385.1 |
| GCF_000014125.1 | GCF_000017865.1 | GCF_000022025.1 | GCF_000025225.2 | GCF_000092985.1 | GCF_000183405.1 |
| GCF_000014145.1 | GCF_000018025.1 | GCF_000022065.1 | GCF_000025265.1 | GCF_000093025.1 | GCF_000183425.1 |
| GCF_000014185.1 | GCF_000018105.1 | GCF_000022085.1 | GCF_000025305.1 | GCF_000093085.1 | GCF_000183725.1 |
| GCF_000014225.1 | GCF_000018145.1 | GCF_000022125.1 | GCF_000025345.1 | GCF_000143085.1 | GCF_000183745.1 |
| GCF_000014265.1 | GCF_000018205.1 | GCF_000022145.1 | GCF_000025405.2 | GCF_000143145.1 | GCF_000184345.1 |
| GCF_000014285.2 | GCF_000018265.1 | GCF_000022265.1 | GCF_000025485.1 | GCF_000143165.1 | GCF_000184435.1 |
| GCF_000014345.1 | GCF_000018285.1 | GCF_000022325.1 | GCF_000025605.1 | GCF_000143845.1 | GCF_000184685.1 |
| GCF_000014385.1 | GCF_000018325.1 | GCF_000022525.1 | GCF_000025645.1 | GCF_000143965.1 | GCF_000184705.1 |

| | | | | | |
|---|---|---|---|---|---|
| GCF_000184745.1 | GCF_000212375.1 | GCF_000236685.1 | GCF_000284015.1 | GCF_000331995.1 | GCF_000512735.1 |
| GCF_000185805.1 | GCF_000212395.1 | GCF_000236705.1 | GCF_000284035.1 | GCF_000332115.1 | GCF_000512895.1 |
| GCF_000185885.1 | GCF_000212415.1 | GCF_000237065.1 | GCF_000284075.1 | GCF_000332735.1 | GCF_000512915.1 |
| GCF_000185965.1 | GCF_000212675.2 | GCF_000237085.1 | GCF_000284095.1 | GCF_000340435.2 | GCF_000513215.1 |
| GCF_000186005.1 | GCF_000212695.1 | GCF_000237205.1 | GCF_000284115.1 | GCF_000340885.1 | GCF_000513295.1 |
| GCF_000186245.1 | GCF_000212735.1 | GCF_000237995.1 | GCF_000284155.1 | GCF_000341345.1 | GCF_000517425.1 |
| GCF_000186265.1 | GCF_000213255.1 | GCF_000238215.1 | GCF_000284255.1 | GCF_000341355.1 | GCF_000517565.1 |
| GCF_000186345.1 | GCF_000213805.1 | GCF_000238255.3 | GCF_000284295.1 | GCF_000341385.1 | GCF_000517605.1 |
| GCF_000186385.1 | GCF_000213825.1 | GCF_000240075.2 | GCF_000284315.1 | GCF_000341395.1 | GCF_000521505.1 |
| GCF_000187005.1 | GCF_000214095.2 | GCF_000240165.1 | GCF_000284335.1 | GCF_000344785.1 | GCF_000521565.1 |
| GCF_000189295.2 | GCF_000214155.1 | GCF_000242255.2 | GCF_000284375.1 | GCF_000344805.1 | GCF_000523235.1 |
| GCF_000189415.1 | GCF_000214175.1 | GCF_000242455.2 | GCF_000284415.1 | GCF_000347595.1 | GCF_000524555.1 |
| GCF_000189535.1 | GCF_000214215.1 | GCF_000242595.2 | GCF_000284515.1 | GCF_000347635.1 | GCF_000525635.1 |
| GCF_000189775.2 | GCF_000214355.1 | GCF_000242635.2 | GCF_000284615.1 | GCF_000347695.1 | GCF_000525655.1 |
| GCF_000190435.1 | GCF_000214375.1 | GCF_000243115.2 | GCF_000286435.2 | GCF_000348725.1 | GCF_000525675.1 |
| GCF_000190535.1 | GCF_000214435.1 | GCF_000243135.2 | GCF_000287335.1 | GCF_000348805.1 | GCF_000550765.1 |
| GCF_000190555.1 | GCF_000214665.1 | GCF_000243155.2 | GCF_000287355.1 | GCF_000349975.1 | GCF_000550785.1 |
| GCF_000190575.1 | GCF_000214785.1 | GCF_000246855.1 | GCF_000294515.1 | GCF_000354175.2 | GCF_000550805.1 |
| GCF_000190595.1 | GCF_000214825.1 | GCF_000247565.1 | GCF_000294775.2 | GCF_000355675.1 | GCF_000565175.1 |
| GCF_000190635.1 | GCF_000215085.1 | GCF_000247605.1 | GCF_000297055.2 | GCF_000355765.4 | GCF_000565195.1 |
| GCF_000190735.1 | GCF_000215105.1 | GCF_000247715.1 | GCF_000298115.2 | GCF_000367205.1 | GCF_000565215.1 |
| GCF_000191045.1 | GCF_000215705.1 | GCF_000248095.2 | GCF_000299235.1 | GCF_000376545.2 | GCF_000568815.1 |
| GCF_000191545.1 | GCF_000215975.1 | GCF_000250635.1 | GCF_000299335.2 | GCF_000376585.1 | GCF_000576555.1 |
| GCF_000192745.1 | GCF_000217635.1 | GCF_000250655.1 | GCF_000299355.1 | GCF_000376645.1 | GCF_000577895.1 |
| GCF_000192865.1 | GCF_000217675.1 | GCF_000250675.2 | GCF_000300005.1 | GCF_000380335.1 | GCF_000583875.1 |
| GCF_000193395.1 | GCF_000217795.1 | GCF_000252445.1 | GCF_000300135.1 | GCF_000385925.1 | GCF_000590475.1 |
| GCF_000194135.1 | GCF_000217815.1 | GCF_000252855.1 | GCF_000300235.2 | GCF_000389635.1 | GCF_000590555.1 |
| GCF_000194605.1 | GCF_000218545.1 | GCF_000253015.1 | GCF_000300295.4 | GCF_000397205.1 | GCF_000590925.1 |
| GCF_000195275.1 | GCF_000218565.1 | GCF_000253035.1 | GCF_000300455.3 | GCF_000400935.1 | GCF_000597865.1 |
| GCF_000195295.1 | GCF_000218625.1 | GCF_000253275.1 | GCF_000304215.1 | GCF_000400955.1 | GCF_000599985.1 |
| GCF_000195315.1 | GCF_000218875.1 | GCF_000253395.1 | GCF_000304455.1 | GCF_000412695.1 | GCF_000600005.1 |
| GCF_000195335.1 | GCF_000218895.1 | GCF_000255115.2 | GCF_000304735.1 | GCF_000418365.1 | GCF_000604125.1 |
| GCF_000195775.1 | GCF_000219105.1 | GCF_000255135.1 | GCF_000305785.2 | GCF_000422085.1 | GCF_000612055.1 |
| GCF_000195875.1 | GCF_000219215.1 | GCF_000255295.1 | GCF_000305935.1 | GCF_000439435.1 | GCF_000612485.1 |
| GCF_000196015.1 | GCF_000219535.2 | GCF_000255535.1 | GCF_000306785.1 | GCF_000439455.1 | GCF_000612685.1 |
| GCF_000196135.1 | GCF_000219605.1 | GCF_000258405.1 | GCF_000306885.1 | GCF_000442645.1 | GCF_000619905.2 |
| GCF_000196175.1 | GCF_000219725.1 | GCF_000259175.1 | GCF_000307105.1 | GCF_000444875.1 | GCF_000626635.1 |
| GCF_000196255.1 | GCF_000219805.1 | GCF_000259255.1 | GCF_000307165.1 | GCF_000444995.1 | GCF_000626675.1 |
| GCF_000196275.1 | GCF_000220625.1 | GCF_000259275.1 | GCF_000309885.1 | GCF_000445425.4 | GCF_000632805.1 |
| GCF_000196295.1 | GCF_000222305.1 | GCF_000260965.1 | GCF_000311765.1 | GCF_000447675.1 | GCF_000632985.1 |
| GCF_000196315.1 | GCF_000222485.1 | GCF_000260985.4 | GCF_000313175.2 | GCF_000454045.1 | GCF_000635915.2 |
| GCF_000196355.1 | GCF_000224005.2 | GCF_000262305.1 | GCF_000313635.1 | GCF_000455605.1 | GCF_000648515.1 |
| GCF_000196395.1 | GCF_000224085.1 | GCF_000264455.2 | GCF_000316515.1 | GCF_000463355.1 | GCF_000661895.1 |
| GCF_000196435.1 | GCF_000224105.1 | GCF_000264765.2 | GCF_000317025.1 | GCF_000463505.1 | GCF_000695095.2 |
| GCF_000196455.1 | GCF_000224985.1 | GCF_000265295.1 | GCF_000317105.1 | GCF_000468615.2 | GCF_000695835.1 |
| GCF_000196495.1 | GCF_000225325.1 | GCF_000265365.1 | GCF_000317125.1 | GCF_000470775.1 | GCF_000696485.1 |
| GCF_000196515.1 | GCF_000225345.1 | GCF_000265385.1 | GCF_000317305.3 | GCF_000471025.2 | GCF_000697965.2 |
| GCF_000196535.1 | GCF_000225445.1 | GCF_000265405.1 | GCF_000317435.1 | GCF_000473245.1 | GCF_000699505.1 |
| GCF_000196615.1 | GCF_000225465.1 | GCF_000265425.1 | GCF_000317475.1 | GCF_000473995.1 | GCF_000706685.1 |
| GCF_000196675.1 | GCF_000226295.1 | GCF_000265465.1 | GCF_000317495.1 | GCF_000477415.1 | GCF_000723165.1 |
| GCF_000196695.1 | GCF_000226315.1 | GCF_000265505.1 | GCF_000317575.1 | GCF_000478885.1 | GCF_000723425.2 |
| GCF_000196735.1 | GCF_000226565.1 | GCF_000266885.1 | GCF_000317615.1 | GCF_000484505.1 | GCF_000723465.1 |
| GCF_000196815.1 | GCF_000226625.1 | GCF_000266905.1 | GCF_000317675.1 | GCF_000484535.1 | GCF_000723505.1 |
| GCF_000196855.1 | GCF_000227665.2 | GCF_000266925.1 | GCF_000317695.1 | GCF_000485905.1 | GCF_000724485.1 |
| GCF_000197735.1 | GCF_000227685.2 | GCF_000266945.1 | GCF_000317835.1 | GCF_000493735.1 | GCF_000724605.1 |
| GCF_000198775.1 | GCF_000227705.2 | GCF_000269925.1 | GCF_000317855.1 | GCF_000494755.1 | GCF_000724625.1 |
| GCF_000199675.1 | GCF_000227745.2 | GCF_000269985.1 | GCF_000317975.2 | GCF_000495505.1 | GCF_000724775.3 |
| GCF_000200595.1 | GCF_000230275.1 | GCF_000270085.1 | GCF_000319245.1 | GCF_000495935.2 | GCF_000725365.1 |
| GCF_000202635.1 | GCF_000230555.1 | GCF_000270245.1 | GCF_000319575.2 | GCF_000497525.1 | GCF_000725405.1 |
| GCF_000202835.1 | GCF_000230695.2 | GCF_000271405.2 | GCF_000321415.2 | GCF_000500935.1 | GCF_000730165.1 |
| GCF_000203895.1 | GCF_000230895.2 | GCF_000271665.2 | GCF_000325665.1 | GCF_000503895.1 | GCF_000730385.1 |
| GCF_000204135.1 | GCF_000231385.2 | GCF_000276685.1 | GCF_000325705.1 | GCF_000504125.1 | GCF_000731315.1 |
| GCF_000204155.1 | GCF_000231405.2 | GCF_000277125.1 | GCF_000325745.1 | GCF_000507245.1 | GCF_000732925.1 |
| GCF_000204255.1 | GCF_000233715.2 | GCF_000277715.1 | GCF_000327045.1 | GCF_000508225.1 | GCF_000732945.1 |
| GCF_000204565.1 | GCF_000233775.1 | GCF_000277795.1 | GCF_000328625.1 | GCF_000508245.1 | GCF_000734015.1 |
| GCF_000204645.1 | GCF_000233915.3 | GCF_000279145.1 | GCF_000328705.1 | GCF_000511305.1 | GCF_000736415.1 |
| GCF_000208385.1 | GCF_000235405.2 | GCF_000281175.1 | GCF_000328725.1 | GCF_000511355.1 | GCF_000737325.1 |
| GCF_000208405.1 | GCF_000235605.1 | GCF_000283575.1 | GCF_000328765.2 | GCF_000511385.1 | GCF_000737865.1 |
| GCF_000209675.1 | GCF_000236405.1 | GCF_000283595.1 | GCF_000330885.1 | GCF_000512205.2 | GCF_000739085.1 |
| GCF_000210915.2 | GCF_000236665.1 | GCF_000283615.1 | GCF_000331735.1 | GCF_000512355.1 | GCF_000739375.1 |

| | | | | | |
|---|---|---|---|---|---|
| GCF_000739435.1 | GCF_000829395.1 | GCF_001017435.1 | GCF_001298465.1 | GCF_001543145.1 | GCF_001636295.1 |
| GCF_000739455.1 | GCF_000830005.1 | GCF_001017655.1 | GCF_001298525.1 | GCF_001543175.1 | GCF_001641005.1 |
| GCF_000739475.1 | GCF_000830885.1 | GCF_001020955.1 | GCF_001302565.1 | GCF_001543245.1 | GCF_001641285.1 |
| GCF_000743945.1 | GCF_000831005.1 | GCF_001020985.1 | GCF_001302585.1 | GCF_001543285.1 | GCF_001642085.1 |
| GCF_000746645.1 | GCF_000831485.1 | GCF_001021025.1 | GCF_001304715.1 | GCF_001543325.1 | GCF_001642655.1 |
| GCF_000747315.1 | GCF_000831645.3 | GCF_001021045.1 | GCF_001305575.2 | GCF_001544015.1 | GCF_001643015.1 |
| GCF_000747585.1 | GCF_000832305.1 | GCF_001021065.1 | GCF_001305595.1 | GCF_001545095.1 | GCF_001643775.1 |
| GCF_000750535.1 | GCF_000832605.1 | GCF_001021935.1 | GCF_001307195.1 | GCF_001545155.1 | GCF_001643955.1 |
| GCF_000754275.1 | GCF_000832905.1 | GCF_001022195.1 | GCF_001307545.1 | GCF_001547735.1 | GCF_001643975.1 |
| GCF_000755145.1 | GCF_000832985.1 | GCF_001025155.1 | GCF_001307805.1 | GCF_001547755.1 | GCF_001644565.1 |
| GCF_000756615.1 | GCF_000833025.1 | GCF_001025175.1 | GCF_001308145.2 | GCF_001547935.1 | GCF_001644605.1 |
| GCF_000757785.1 | GCF_000833105.2 | GCF_001026985.1 | GCF_001308265.1 | GCF_001547975.1 | GCF_001644705.1 |
| GCF_000757825.1 | GCF_000833575.1 | GCF_001027025.1 | GCF_001308285.1 | GCF_001547995.1 | GCF_001647635.1 |
| GCF_000758665.1 | GCF_000835165.1 | GCF_001027285.1 | GCF_001310085.1 | GCF_001548015.1 | GCF_001648115.1 |
| GCF_000758685.1 | GCF_000875755.1 | GCF_001027545.1 | GCF_001310225.1 | GCF_001548055.1 | GCF_001648175.1 |
| GCF_000758725.1 | GCF_000935025.1 | GCF_001028625.1 | GCF_001314245.2 | GCF_001548275.1 | GCF_001652465.1 |
| GCF_000759475.1 | GCF_000940805.1 | GCF_001028645.1 | GCF_001314305.1 | GCF_001553195.1 | GCF_001652485.1 |
| GCF_000761155.1 | GCF_000940845.1 | GCF_001028705.1 | GCF_001314325.1 | GCF_001553545.1 | GCF_001653335.1 |
| GCF_000761215.1 | GCF_000940995.1 | GCF_001029265.1 | GCF_001314945.1 | GCF_001553565.1 | GCF_001653755.1 |
| GCF_000763535.1 | GCF_000941055.1 | GCF_001038625.1 | GCF_001314995.1 | GCF_001553605.1 | GCF_001653955.1 |
| GCF_000763575.1 | GCF_000941075.1 | GCF_001040945.1 | GCF_001315015.1 | GCF_001553625.1 | GCF_001654435.1 |
| GCF_000764535.1 | GCF_000953135.1 | GCF_001042405.1 | GCF_001318345.1 | GCF_001553935.1 | GCF_001654455.1 |
| GCF_000767055.1 | GCF_000953195.1 | GCF_001042595.1 | GCF_001399775.1 | GCF_001553955.1 | GCF_001655245.1 |
| GCF_000767465.1 | GCF_000953215.1 | GCF_001042635.1 | GCF_001402875.1 | GCF_001558255.2 | GCF_001659785.1 |
| GCF_000767615.3 | GCF_000953355.1 | GCF_001042675.1 | GCF_001411495.1 | GCF_001558415.2 | GCF_001660005.1 |
| GCF_000772105.1 | GCF_000953475.1 | GCF_001042695.1 | GCF_001414055.1 | GCF_001559015.1 | GCF_001660485.1 |
| GCF_000785105.2 | GCF_000953635.1 | GCF_001042715.1 | GCF_001420915.1 | GCF_001559115.2 | GCF_001661675.2 |
| GCF_000785495.1 | GCF_000953655.1 | GCF_001043175.1 | GCF_001421015.2 | GCF_001561955.1 | GCF_001663155.1 |
| GCF_000785515.1 | GCF_000953695.1 | GCF_001046955.1 | GCF_001431725.1 | GCF_001562115.1 | GCF_001663175.1 |
| GCF_000785705.2 | GCF_000953715.1 | GCF_001050115.1 | GCF_001432245.1 | GCF_001563225.1 | GCF_001663675.1 |
| GCF_000789395.1 | GCF_000953735.1 | GCF_001050435.1 | GCF_001441165.1 | GCF_001564455.1 | GCF_001676705.1 |
| GCF_000800295.1 | GCF_000954135.1 | GCF_001050475.1 | GCF_001442535.1 | GCF_001572725.1 | GCF_001676785.2 |
| GCF_000800395.1 | GCF_000959245.1 | GCF_001051995.2 | GCF_001442785.1 | GCF_001577305.1 | GCF_001677275.1 |
| GCF_000800475.2 | GCF_000959725.1 | GCF_001077715.1 | GCF_001443605.1 | GCF_001578185.1 | GCF_001677435.1 |
| GCF_000801275.2 | GCF_000960975.1 | GCF_001077815.2 | GCF_001443625.1 | GCF_001578205.1 | GCF_001678905.1 |
| GCF_000801295.1 | GCF_000961095.1 | GCF_001078275.1 | GCF_001444405.1 | GCF_001579945.1 | GCF_001678945.1 |
| GCF_000802245.2 | GCF_000963865.1 | GCF_001182745.1 | GCF_001444425.1 | GCF_001580455.1 | GCF_001679725.1 |
| GCF_000803645.1 | GCF_000964565.1 | GCF_001187505.1 | GCF_001444445.1 | GCF_001583415.1 | GCF_001682385.1 |
| GCF_000807255.1 | GCF_000967305.2 | GCF_001187595.1 | GCF_001445575.1 | GCF_001584185.1 | GCF_001685355.1 |
| GCF_000807275.1 | GCF_000967895.1 | GCF_001187785.1 | GCF_001447335.1 | GCF_001584725.1 | GCF_001685395.1 |
| GCF_000807675.2 | GCF_000967915.1 | GCF_001187845.1 | GCF_001454945.1 | GCF_001586165.1 | GCF_001685435.2 |
| GCF_000812665.2 | GCF_000968055.1 | GCF_001189295.1 | GCF_001455205.1 | GCF_001586195.1 | GCF_001685465.1 |
| GCF_000815025.1 | GCF_000968195.1 | GCF_001189495.1 | GCF_001456065.2 | GCF_001586215.1 | GCF_001687475.2 |
| GCF_000815065.1 | GCF_000968535.2 | GCF_001190745.1 | GCF_001456115.1 | GCF_001586255.1 | GCF_001687545.1 |
| GCF_000815185.1 | GCF_000969765.1 | GCF_001190755.1 | GCF_001457455.1 | GCF_001590605.1 | GCF_001687565.2 |
| GCF_000815225.1 | GCF_000972865.1 | GCF_001190945.1 | GCF_001457475.1 | GCF_001590685.1 | GCF_001688625.2 |
| GCF_000816085.1 | GCF_000973085.1 | GCF_001262015.1 | GCF_001460635.1 | GCF_001594265.1 | GCF_001693385.1 |
| GCF_000816145.1 | GCF_000973105.1 | GCF_001262055.1 | GCF_001465255.1 | GCF_001597285.1 | GCF_001693675.1 |
| GCF_000816185.1 | GCF_000973505.1 | GCF_001262715.1 | GCF_001465295.1 | GCF_001598035.1 | GCF_001697185.1 |
| GCF_000816345.1 | GCF_000973545.1 | GCF_001263175.1 | GCF_001465795.2 | GCF_001602095.1 | GCF_001697225.1 |
| GCF_000816845.1 | GCF_000973625.1 | GCF_001263395.1 | GCF_001465835.2 | GCF_001605015.1 | GCF_001698145.1 |
| GCF_000817955.1 | GCF_000973725.1 | GCF_001267155.1 | GCF_001465855.1 | GCF_001606005.1 | GCF_001698205.1 |
| GCF_000817975.1 | GCF_000974425.1 | GCF_001267175.1 | GCF_001466725.1 | GCF_001606025.1 | GCF_001698225.1 |
| GCF_000818015.1 | GCF_000974685.2 | GCF_001267925.1 | GCF_001482365.1 | GCF_001610955.1 | GCF_001700895.1 |
| GCF_000818035.1 | GCF_000980815.1 | GCF_001273775.1 | GCF_001482385.1 | GCF_001610975.1 | GCF_001700965.1 |
| GCF_000818095.1 | GCF_000980835.1 | GCF_001273795.1 | GCF_001483385.1 | GCF_001611135.1 | GCF_001701025.1 |
| GCF_000819445.1 | GCF_000981525.1 | GCF_001274875.1 | GCF_001483865.1 | GCF_001611675.1 | GCF_001701045.1 |
| GCF_000819565.1 | GCF_000981765.1 | GCF_001274895.1 | GCF_001484065.1 | GCF_001611975.1 | GCF_001702135.1 |
| GCF_000827005.1 | GCF_000982715.1 | GCF_001275345.1 | GCF_001484605.1 | GCF_001617605.1 | GCF_001702175.1 |
| GCF_000827125.1 | GCF_000982825.1 | GCF_001277995.1 | GCF_001484935.1 | GCF_001617625.1 | GCF_001704115.1 |
| GCF_000828475.1 | GCF_000987835.1 | GCF_001278055.1 | GCF_001499615.1 | GCF_001618885.1 | GCF_001704155.1 |
| GCF_000828615.1 | GCF_000993785.2 | GCF_001278075.1 | GCF_001499655.1 | GCF_001620265.1 | GCF_001704275.1 |
| GCF_000828635.1 | GCF_001005905.1 | GCF_001280225.1 | GCF_001507645.1 | GCF_001620305.1 | GCF_001704615.2 |
| GCF_000828655.1 | GCF_001006005.1 | GCF_001281045.1 | GCF_001509405.1 | GCF_001623565.1 | GCF_001705565.1 |
| GCF_000828675.1 | GCF_001007875.1 | GCF_001281405.1 | GCF_001517405.1 | GCF_001629705.1 | GCF_001708305.1 |
| GCF_000828835.1 | GCF_001007935.1 | GCF_001281465.1 | GCF_001534665.1 | GCF_001632775.1 | GCF_001708405.1 |
| GCF_000828855.1 | GCF_001007995.1 | GCF_001281485.1 | GCF_001534745.1 | GCF_001632805.1 | GCF_001709315.1 |
| GCF_000829035.1 | GCF_001008165.2 | GCF_001293125.1 | GCF_001542625.1 | GCF_001633165.1 | GCF_001715535.1 |
| GCF_000829235.1 | GCF_001010285.1 | GCF_001294625.1 | GCF_001542775.1 | GCF_001634285.1 | GCF_001717505.1 |
| GCF_000829315.1 | GCF_001013905.1 | GCF_001295365.1 | GCF_001543105.1 | GCF_001636015.1 | GCF_001717955.1 |

| | | | |
|---|---|---|---|
| GCF_001719165.1 | GCF_001941565.1 | GCF_002080435.1 | GCF_900087055.1 |
| GCF_001720485.1 | GCF_001941805.1 | GCF_002082015.1 | GCF_900089455.2 |
| GCF_001721185.1 | GCF_001941825.1 | GCF_002082605.1 | GCF_900090215.1 |
| GCF_001721645.1 | GCF_001941945.1 | GCF_002096055.1 | GCF_900093775.1 |
| GCF_001721685.1 | GCF_001945665.1 | GCF_002097535.1 | GCF_900095135.1 |
| GCF_001729245.1 | GCF_001951155.1 | GCF_002101335.1 | GCF_900095795.1 |
| GCF_001729485.1 | GCF_001951175.1 | GCF_002101395.1 | GCF_900097105.1 |
| GCF_001729525.1 | GCF_001953055.1 | GCF_002104335.1 | GCF_900116045.1 |
| GCF_001729725.1 | GCF_001953075.1 | GCF_002105555.1 | GCF_900116935.1 |
| GCF_001735765.2 | GCF_001953175.1 | GCF_002105755.1 | GCF_900120165.1 |
| GCF_001735805.1 | GCF_001953195.1 | GCF_002109385.1 | GCF_900120345.1 |
| GCF_001741865.1 | GCF_001953955.1 | GCF_002116905.1 | GCF_900120375.1 |
| GCF_001742225.1 | GCF_001955715.1 | GCF_002117085.1 | GCF_900128415.1 |
| GCF_001746835.1 | GCF_001955735.1 | GCF_002117105.1 | GCF_900128595.1 |
| GCF_001747405.1 | GCF_001956985.1 | GCF_002117405.1 | GCF_900128725.1 |
| GCF_001747425.1 | GCF_001969365.1 | GCF_002117445.1 | GCF_900128735.1 |
| GCF_001750165.1 | GCF_001969385.1 | GCF_002119665.1 | GCF_900155405.1 |
| GCF_001750685.1 | GCF_001971565.1 | GCF_002119765.1 | GCF_900155415.1 |
| GCF_001753205.1 | GCF_001971745.1 | GCF_002119805.1 | GCF_900168365.1 |
| GCF_001753245.1 | GCF_001974985.1 | GCF_002127965.1 | GCF_900169085.1 |
| GCF_001761345.1 | GCF_001975025.1 | GCF_002142495.1 | GCF_900169485.1 |
| GCF_001761365.1 | GCF_001975225.1 | GCF_002142615.1 | GCF_900169565.1 |
| GCF_001761465.1 | GCF_001975665.1 | GCF_002157205.1 | |
| GCF_001766235.1 | GCF_001975705.1 | GCF_002157855.1 | |
| GCF_001767235.1 | GCF_001983935.1 | GCF_002157895.1 | |
| GCF_001767275.1 | GCF_001983975.1 | GCF_002158865.1 | |
| GCF_001787335.1 | GCF_001983995.1 | GCF_002158905.1 | |
| GCF_001831475.1 | GCF_001988955.1 | GCF_002162335.1 | |
| GCF_001831495.1 | GCF_001989575.1 | GCF_002162375.1 | |
| GCF_001854125.1 | GCF_001997295.1 | GCF_002163585.1 | |
| GCF_001854185.1 | GCF_001998765.1 | GCF_002163975.1 | |
| GCF_001854245.1 | GCF_001998865.1 | GCF_002173495.1 | |
| GCF_001858005.1 | GCF_001998885.1 | GCF_002189675.2 | |
| GCF_001865575.1 | GCF_001999245.1 | GCF_002192415.1 | |
| GCF_001865855.1 | GCF_001999905.1 | GCF_002192455.1 | |
| GCF_001870205.1 | GCF_001999945.1 | GCF_002196515.1 | |
| GCF_001870665.2 | GCF_001999985.1 | GCF_002201795.1 | |
| GCF_001877035.1 | GCF_002000985.1 | GCF_002208135.1 | |
| GCF_001878675.1 | GCF_002003265.1 | GCF_002208805.2 | |
| GCF_001880325.1 | GCF_002005145.1 | GCF_002208825.2 | |
| GCF_001886615.1 | GCF_002005165.1 | GCF_002209165.2 | |
| GCF_001886695.1 | GCF_002005305.1 | GCF_002209185.2 | |
| GCF_001886715.1 | GCF_002005365.1 | GCF_002209385.1 | |
| GCF_001886815.1 | GCF_002005405.1 | GCF_002211765.1 | |
| GCF_001886855.1 | GCF_002005445.1 | GCF_002211785.1 | |
| GCF_001887245.1 | GCF_002005465.1 | GCF_002214645.1 | |
| GCF_001887285.1 | GCF_002006565.1 | GCF_002215135.1 | |
| GCF_001888165.1 | GCF_002007565.1 | GCF_002215215.1 | |
| GCF_001888185.1 | GCF_002007745.1 | GCF_002218195.1 | |
| GCF_001889165.1 | GCF_002009175.1 | GCF_002218245.1 | |
| GCF_001900245.1 | GCF_002021925.1 | GCF_002220735.1 | |
| GCF_001902315.1 | GCF_002024185.1 | GCF_002221525.1 | |
| GCF_001908775.1 | GCF_002024265.1 | GCF_002222615.2 | |
| GCF_001913135.1 | GCF_002025645.1 | GCF_002222635.1 | |
| GCF_001922025.1 | GCF_002025665.1 | GCF_002222655.1 | |
| GCF_001922305.1 | GCF_002025725.1 | GCF_002224365.1 | |
| GCF_001922385.1 | GCF_002028405.1 | GCF_002224425.1 | |
| GCF_001922545.1 | GCF_002043005.1 | GCF_002224565.1 | |
| GCF_001931675.1 | GCF_002056725.1 | GCF_002224645.1 | |
| GCF_001932615.1 | GCF_002056795.1 | GCF_002234495.1 | |
| GCF_001936235.1 | GCF_002067135.1 | GCF_002234535.1 | |
| GCF_001936255.1 | GCF_002073635.2 | GCF_002237615.1 | |
| GCF_001936335.1 | GCF_002073715.2 | GCF_002240035.1 | |
| GCF_001940525.2 | GCF_002074155.1 | GCF_002240355.1 | |
| GCF_001941345.1 | GCF_002075105.1 | GCF_002240375.1 | |
| GCF_001941425.1 | GCF_002075285.2 | GCF_900016785.1 | |
| GCF_001941445.1 | GCF_002075795.1 | GCF_900070355.1 | |
| GCF_001941465.1 | GCF_002079945.1 | GCF_900078695.1 | |
| GCF_001941485.1 | GCF_002080395.1 | GCF_900078775.1 | |
| GCF_001941505.1 | GCF_002080415.1 | GCF_900086555.1 | |
